# Spatiotemporal atlas of the fetal brain depicts cortical developmental gradient in Chinese population

**DOI:** 10.1101/2022.05.09.491258

**Authors:** Xinyi Xu, Cong Sun, Jiwei Sun, Wen Shi, Yao Shen, Ruoke Zhao, Wanrong Luo, Mingyang Li, Guangbin Wang, Dan Wu

## Abstract

The fetal brains experience rapid and complex development in utero during the second and third trimesters. MRI atlas of the fetal brain in this period enables us to quantify the normal fetal brain development in the spatiotemporal domain. In this study, we constructed a high quality spatiotemporal atlas between 23-38 weeks gestational age (GA) from 90 healthy Chinese fetuses using a pairwise and groupwise registration-based pipeline. We quantified the fetal cortical morphology and characterized the developmental trajectories of each morphological index. The cortical thickness exhibited a biphasic growth pattern; the developmental trend of curvature fitted well into the Gompertz model; sulcal depth increased linearly while surface area expanded exponentially. The cortical thickness and curvature trajectories consistently pointed to a characteristic time-point during development around 31 weeks of GA. The characteristic GA and growth rate obtained from individual cortical regions suggested a central-to-peripheral developmental gradient, with the earliest development in the parietal lobe, and we also observed within-lobe regional orders, which may be linked to biophysical events such as dendritic arborization and thalamocortical fibers ingrowth. The proposed atlas was also compared with an existing fetal atlas from a Caucasian/mixed population. Finally, we examined the structural asymmetry of the fetal brains and found extensive asymmetry that dynamically changed with development. The current study depicted a comprehensive profile of fetal cortical development and the established atlas could be used as a normative reference for neurodevelopmental and diagnostic purposes, especially in the Chinese population.

## 1 Introduction

The cerebrum of fetuses undergoes complex yet well-orchestrated development during gestation, especially in the cerebral cortex. There are a range of histogenetic processes involved in cortical development, including neuronal migration, proliferation, dendritic differentiation, axonal ingrowth and outgrowth^1,2^. The developmental trajectory of normal fetal cerebral cortex may reflect the underlying cellular processes, yet due to the lack of *in vivo* examination tools, details of the cortical development are not well elucidated. At present, in-utero MRI has become one of the widely adopted medical imaging tools for prenatal examination given its rich contrast and superior resolution compared to ultrasound. Despite the challenges in fetal brain MRI, which is subjected to fetal and maternal motion, advanced acquisition and post-processing technologies have been developed in the past decade to make it feasible for 3D visualization of the fetal brain ^3–16^.

Developing atlas of the healthy fetal brain provides a standard anatomical reference at each gestational age (GA), which is essential for quantitative image analysis. Habas et al.^17^ developed the first fetal brain atlas using an elastic deformation model and groupwise registration in a narrow GA range of 21 and 24 weeks. Serag et al.^18^ used the free form deformation in combination with an adaptive kernel regression method to generate a fetal brain atlas covering 22 to 38 weeks of GA (brain-development.org), although at relatively low resolution of 1.18 mm isotropic. Gholipour et al.^19^ constructed an unbiased sharp deformable fetal brain atlas using Symmetric diffeomorphic deformation in fetuses between 19-39 weeks of GA (http://crl.med.harvard.edu/research/fetal_brain_atlas/). There are several other fetal brain atlases, such as the postmortem fetal brain atlas from 15 to 22 weeks^10^, the cortical surface atlases of healthy fetuses^11,20^, and craniofacial features reserved fetal brain atlas in the GA range of 21 to 36 weeks^21^. See our summary in Supplementary Table S1.

The aforementioned fetal brain atlases (except for the postmortem atlas^10^) were mostly generated from Caucasian or mixed populations, and it is known that ethnic difference contributes to brain morphology. For instance, Tang et al. reported regional brain volumes differed between Chinese and Caucasian populations^22^. Zhao et al. built a Chinese pediatric brain atlases between 6-12 years old and found dramatic anatomical differences compared to the National Institutes of Health pediatric atlases^23^. Since both genetic and environmental factors could lead to populational differences, investigation in the early developing brain, especially the in-utero fetal brain may help to filter the postnatal factors. Preliminary attempts have been made to establish fetal brain atlases based on the Chinese population^12,24^. However, these two atlases both used 1.5T MRI data that are collected using clinical protocols with thick slices in an limited age range (23 to 36 weeks in every 2 weeks in Li et al.^24^ and 22 to 35 weeks in Wu et al.^12^), making them difficult for direct comparison with the existing 3T atlases. In this study, we aimed to establish an unbiased high quality Chinese fetal atlas covering the second-to-third gestation (23 to 38 weeks GA), using a carefully designed atlas generation pipeline.

Based on the fetal brain atlas, the spatiotemporal developmental pattern can be depicted *in utero*, e.g., curvature, thickness, surface area, and volume in different cortical regions. Previous groups have demonstrated the feasibility of extracting these cortical morphological indicators and showed how they change with GA in selected brain regions, without parameterizing the trajectories. Rajagopalan et al.^5^ characterized developmental trajectories of overall cerebral volume and mean Jacobian determinants in central sulcus, frontal white matter, etc. Several studies have depicted the changes of regional volumes with GA^6,7,12,13^. Wright et al. studied the fetal brain cortical folding pattern with curvature-based indicators^8^. Xia et al. depicted the regional surface areas in 26-29w fetal brains for cortical parcellation^11^. Xu et al. portrayed the cortical developmental trajectories in second trimester postmortem fetal brains^9^. Habas et al. depicted the curvature in different sulci and gyri of 20-28w fetuses^4^. Clouchoux et al. studied the gyrification index and surface area of normal fetuses and fetuses with congenital heart disease^14,15^. Sulcal pattern analysis studies showed that the sulcal differences between abnormal fetuses (e.g., agenesis of corpus callosum, congenital heart disease) and healthy fetuses were detected early in gestation^25,26^. However, a quantitative and comprehensive characterization of the cortical development remains absent, and most of these studies did not link the growth patterns with underlying biological events.

In this work, we proposed a spatiotemporal atlas of the Chinese fetal brain based on 3T high resolution in-utero MRI of healthy fetuses scanned at 22.57 to 39.00 weeks of GA, and compared it with existing atlas from Caucasian/mixed populations. We further characterized the cortical morphological trajectories to quantify the developmental order and rate among different cortical regions that gave the gradient of cortical development. Furthermore, we demonstrated the asymmetry of fetal cerebral cortex in different functional regions.

## 2 Material and Methods

### 2.1 Subjects

We collected 219 fetuses of 20-40 weeks of GA in healthy pregnancy at the Shandong Provincial Hospital from October 2019 to September 2021. Ethical approval was obtained from the Institutional Review Board at the local hospital. Written informed consent was obtained from the participants. Exclusion criteria included visible fetal or maternal motion artifacts on MRI and radiological abnormalities of the fetal brain (e.g., ventriculomegaly and asymmetrical ventricles). The image quality was inspected by experienced radiologists and physicists (S.C. and W.G.B.). At last, 90 normal datasets in the GA range of 22.57 to 39.00 weeks with optimal image quality were included for atlas generation (Fig. 1a). GA distribution of the 90 subjects is shown in Fig. 1b.

**Fig. 1.**
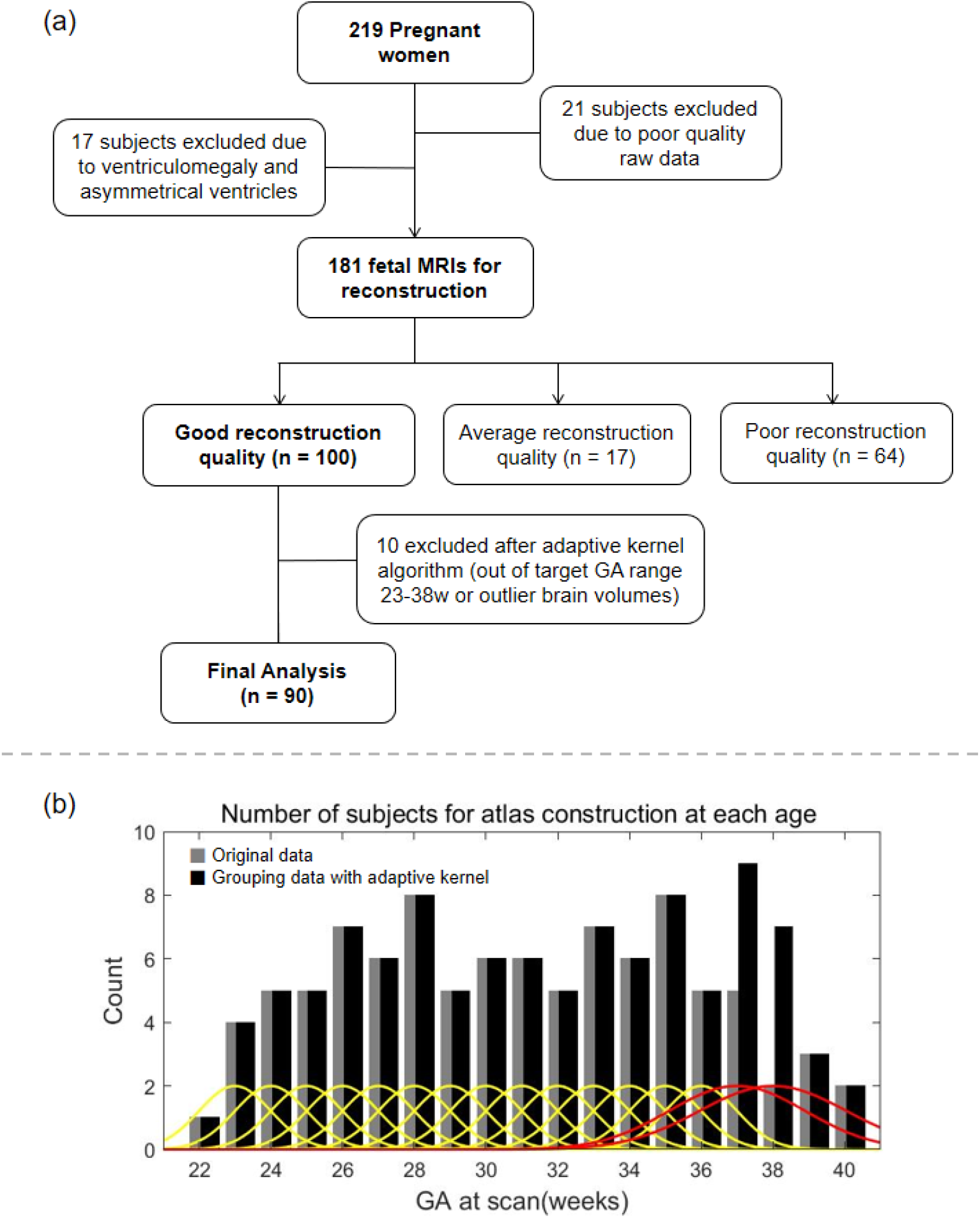
(a) Data inclusion and exclusion flowchart. (b) The number of subjects used for atlas generation in each GA week, determined using an adaptive kernel approach with the weights shown in yellow (regular kernel width for group with a sufficient number of subjects) and red (increased kernel width for group with a small number of subjects).

### 2.2 Image acquisition

The fetal brain MRI was performed on 3.0 T Siemens scanner (MAGNETOM Skyra, Siemens Healthcare, Erlangen, Germany) with an 18-channel body coil. The scans were acquired using T2-weighted Half-Fourier Acquisition Single-shot Turbo spin Echo imaging (HASTE) in multiple orthogonal planes of the fetal brain separately, typically 1 to 4 scans in axial, coronal and sagittal orientations respectively. The imaging parameters were: repetition time = 800 ms, echo time = 113 ms, slice thickness = 2 mm, no slice gap, in-plane resolution = 1.1×1.1 mm^2^, field of view = 280×280 mm^2^, partial Fourier factor = 5/8. No maternal sedation was used.

### 2.3 Image reconstruction

The 2D multislice stacks from multiple orientations went through for preprocessing pipeline to obtain the final 3D volumes, including stack selection, brain extraction, bias field correction, slice-to-volume registration (SVR) and super resolution reconstruction (SRR) (see pipeline in Supplementary Fig. S1). First, 3 to 7 2D stacks with minimal inter-slice motion were manually selected. Then, a Convolutional Neural Network (CNN) based brain extraction tool^27^ was used to automatically extract the fetal brain from each input stacks, followed by an N4 bias field correction algorithm^28^ to correct intensity inhomogeneity. Finally, the 3D high-resolution fetal brain volume was reconstructed at 0.8 mm isotropic resolution using the NiftyMIC^27^ (https://github.com/gift-surg/NiftyMIC.) pipeline, with a two-step iterative SVR-SRR reconstruction. The reconstructed high-resolution volume was rigidly registered to the standard space using FLIRT^29^ for atlas generation.

### 2.4 Atlas Generation

#### Adaptive kernel

In our study, the distribution of subjects over GA was uneven (Fig. 1b histogram in gray). Thus, we performed an adaptive kernel algorithm^30^ to re-group the subjects according to their GA, such that each week contained 5-9 subjects (Fig. 1b histogram in black), except for the 23w group. The weights of each subject were determined by a Gaussian kernel according to the GA:

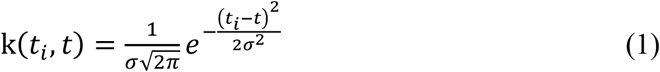

where *i*=*1,2,…, N, N* was the number of subjects at given GA group *t*. *t_i_* denotes GA of the subject *i* at scan. The kernel width *σ* was adjusted according to the number of subjects, i.e., *σ* was iteratively increased (or decreased) when the number of grouped subjects was below (or above) a given number range (e.g., median±tolerance). Here, 0.9*k_max_* was taken as the inclusion threshold, where *k_max_* represents the weights of the subject *i* whose *t_i_* was closest to the target *t*. Note that the image quality of the originally grouped 5 individuals for 37w atlas was relatively low among all the included subjects, so we manually increase the kernel width *σ* to include more subjects. The GA-dependent weights were shown in Fig. 1b in yellow and red.

#### Registration pipeline

We designed a group-wise registration pipeline with pairwise initialization to generate atlas at each GA. We chose symmetric diffeomorphic deformable registration algorithm (SyN)^31^ in which source and target images are affected equally, such that the asymmetric bias can be reduced to preserve unbiased mean anatomy, and it has shown to be more suitable for the developing brains^32^. The atlas generation pipeline, as shown in Fig. 2, can be divided into two steps: initial template construction and iterative update.

**Fig. 2.**
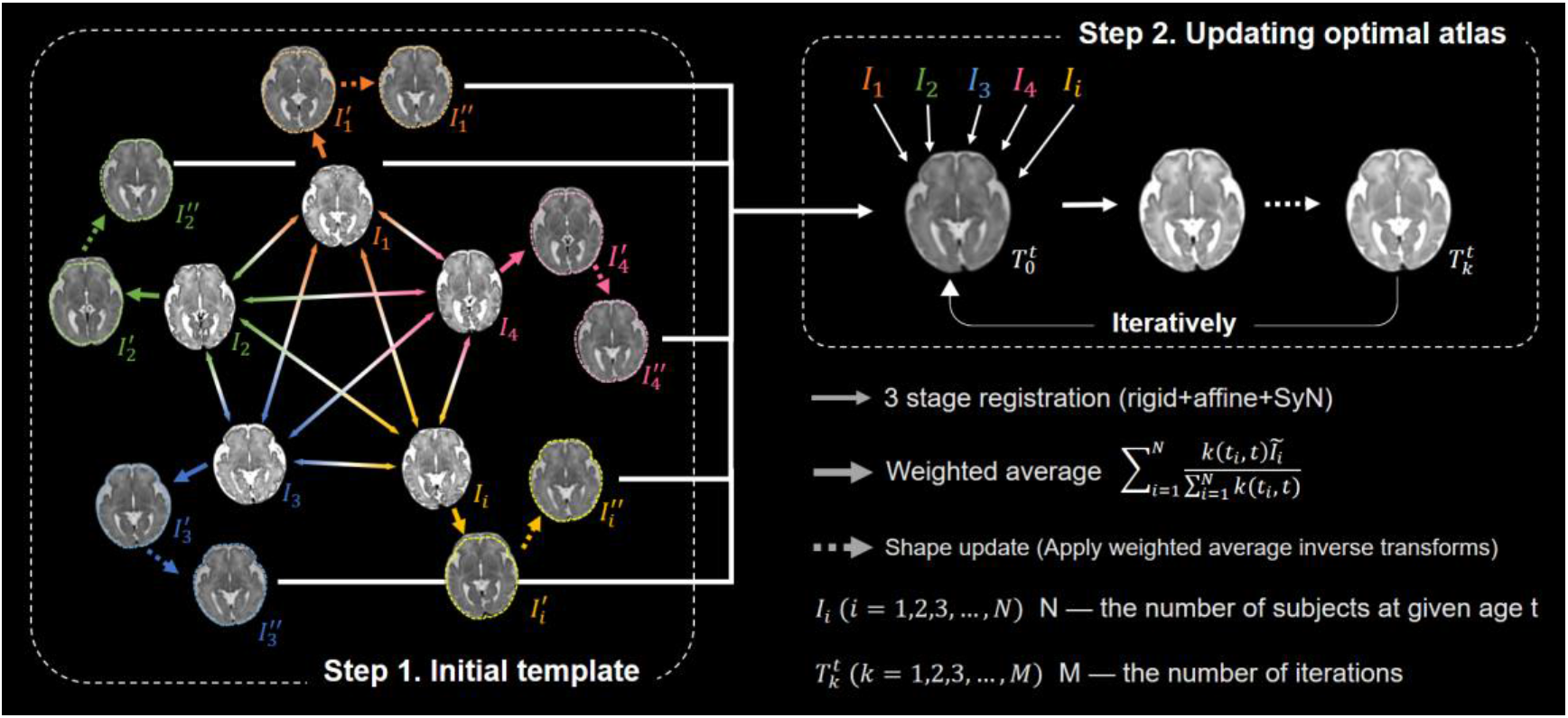
Fetal atlas generation pipeline, including step 1, building the initial template 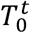, and step 2, iteratively update of the atlas 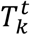. Annotations: *I_i_*, input T2w brain images of subject *i*; 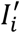, weighted averaged images after pairwise registration; 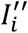, subject images after shape update. Thin arrows represent the rigid, affine, SyN registration in ANTs. Thick arrows represent the weighted average process. Dotted arrows show the shape update procedure. The dashed contours on 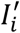 demonstrate the difference in shape before and after shape update.

During the first step, pairwise registration^30,33^ was adopted, where each subject image *I_i_*(*i* = 1,2,…,*N*) was selected as the target in turn, and the rest of subject images were registered to the target using rigid, affine, and SyN registrations in ANTs (https://github.com/ANTsX/ANTs) to obtain the weighted averaged target 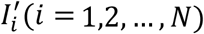. Subsequently, a shape-update procedure^34^ was performed, by applying the weighted averaged inverse transformations on 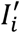 to obtain an unbiased mean image 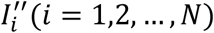, where the weights were obtained according the adaptive kernel k(*t_i_*, *t*) in equation (1). The shape-update is necessary for eliminating bias in the atlas toward any of the individual images (see difference labeled by dashed contours on 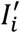 in Fig. 2). The initial template 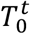 at GA of *t* weeks was constructed by weighted averaging the pairwise registered images 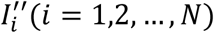.

The second step updated the template iteratively based on group-wise registration. In each iteration *k*, each subject image *I_i_*(*i* = 1,2,…,*N*) was registered to the reference image 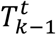 generated during the last iteration, and 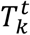 was updated by the weighted average image and shape-update procedure. The second step usually converged in 6 to 10 iterations (Supplementary Fig. S2), and we took the 10^th^ iteration result as our final template at each GA week.

### 2.5 Cortical Analysis

#### Brain segmentation

All 3D reconstructed fetal brain images were segmented using an automatic segmentation DrawEM algorithm (https://github.com/MIRTK/DrawEM)^35^. The cortical gray matter (cGM) label was manually corrected, which was parcellated to 16 regions-of-interest (ROIs) per hemisphere for the following analysis, including the frontal lobe, parietal lobe, occipital lobe, insula, 2 parts in cingulate, and 10 subregions in the temporal lobe.

#### Cortical surface extraction

Based on the refined cortical segmentation, the white matter (WM) surface between cGM and WM labels and pial surface between cGM and CSF labels were reconstructed using dHCP-structural-pipeline^36^ (https://github.com/BioMedIA/dhcp-structural-pipeline), which was modified to adapt to our fetal data (originally defined for neonates). Besides, the inflated surface, very inflated surface, midthickness surface, and sphere surface were generated for calculating the cortical morphological indices. Cortical features including cortical thickness, curvature, sulcal depth, and surface area were generated from 32 cGM ROIs that were propagated from volumetric space to the surface space. In brief, cortical thickness was calculated as Euclidean distance between WM surface and pial surface; curvature was estimated on WM surface and indicated the degree of cortical folding; sulcal depth represented mean convexity or concavity of cortical surface that was estimated during surface inflation; and surface area was measured by summing up the area of all the triangle meshes within the pial surface.

#### Model fitting

The different cortical morphometric indices demonstrated distinct developmental trajectories and were quantified using different models according to the whole-brain averaged trajectory pattern and the goodness-of-fit. The Beta growth and decay model^37^ was chosen to fit the cortical thickness that exhibited a biphasic change with GA. The model is defined as:

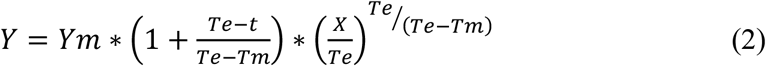

where *Ym* indicates the cortical thickness at peak, and *Te* is the GA when cortical thickness reaches the peak. Mean curvature data were fitted with the commonly used Gompertz growth model^8,38^:

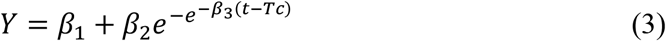

where *β_3_* controls the growth rate, and *Tc* determines the GA where peak growth of curvature occurs. The surface area was fitted using an exponential growth model:

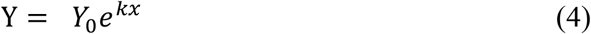

where *k* is the growth rate. The sulcal depth was fitted linearly.

### 2.6 Asymmetry analysis

We divided all the subjects into two GA groups of 23-30w and 31-38w, given the critical changes observed around 31w. Vertex-wise indicators were averaged over different ROIs in left and right hemispheres. Lateralization Index (LI) of each cortical indicator as:

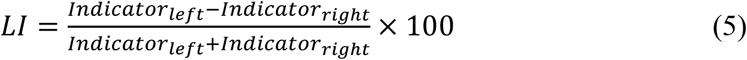

A positive LI in curvature, surface area, and sulcal depth represented the leftward lateralization while negative LI represented the rightward lateralization. Note the lateralization of cortical thickness was not analyzed given the biphasic change. Significance of the asymmetry was calculated using pairwise t-test which was performed on the cortical indicators between left and right ROIs within the group. The p-values were adjusted using the false discovery rate (FDR) method and adjusted p < 0.05 was considered significant.

### 2.7 Difference between CRL atlas and Our atlas

In order to evaluate the fetal brain developmental differences between our Chinese fetal brain atlas (CHN) and CRL atlas based on a Caucasian/mixed population, we obtained the cortical indicators from the CRL atlas data using the same methods as mentioned in section 2.5. Besides, we performed tensor-based morphometry (TBM)^39^ to measure the morphological differences between two atlases at the same GA. We registered the CHN atlas to CRL atlas at corresponding GAs with a 3-stage registration in ANTs including rigid, affine, and SyN, and then calculated the Jacobian determinants from the nonlinear transformation matrix generated during the SyN stage. The deformation vectors were displayed to visualize the local morphological differences between CHN and CRL atlas in terms of direction and amplitude.

## 3 Results

### 3.1 Spatiotemporal atlas of the fetal brain based on Chinese population

The reconstructed CHN atlas (Fig. 3a-c) revealed drastic developmental changes not only in size but also in anatomy, especially in the cortex, e.g., the cortical folding developed rapidly during the second-third trimester. Sylvian fissure (green arrows in Fig. 3 and Fig. S6) was well visualized at 23w of GA, indicating that it was formed before 23w GA ^9^. The central sulcus (yellow arrows in Fig. 3) can be distinguished as early as 24w GA, and it was evident on the curvature maps (Fig. S6). Between 26w to 28w GA, precentral gyrus and postcentral gyrus (orange arrows in Fig. 3 and Fig. S6) gradually emerged. These observations of sulcal development are consistent with the previous studies^4,15,40^. Fig. S6a-c demonstrates the vertex-wise cortical morphological indices on the inflated surface of the proposed CHN atlas at each GA. The CHN atlas is deposited on https://github.com/Thea-Eddie-Amy/CHN-fetal-brain-atlas.

**Fig. 3.**
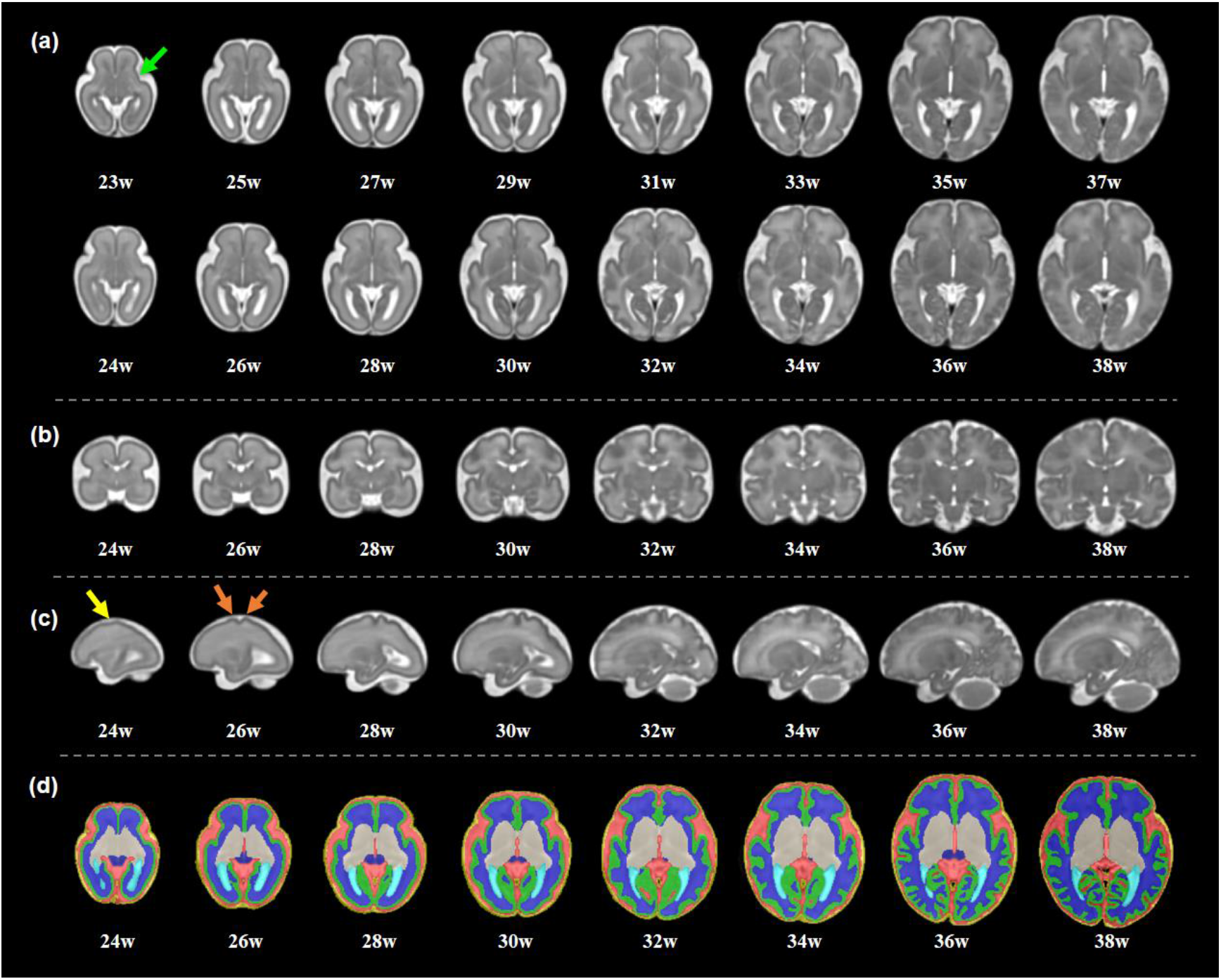
Spatiotemporal structural atlas of fetal brains based on Chinese population. (a) Axial view, (b) coronal view, (c) sagittal view, (d) tissue segmentation after manual correction at representative GAs. Green arrow: Sylvian fissure; yellow arrow: central sulcus; orange arrows: precentral and postcentral gyri.

### 3.2 Cortical morphological developmental trajectories

#### Biphasic growth pattern in cortical thickness

Cortical thickness followed a biphasic change (first increase then decrease) over GA (Fig. 4), and thus, we chose Beta growth and decay model^37^ to fit this pattern. The *Te* parameter of the whole-brain averaged trajectory indicated the peak thickness occurred around 31 weeks GA (Fig. 4b). The *Te* time varied among different regions with the earliest *Te* in the parietal lobe, followed by that in the occipital lobe, temporal lobe, and frontal lobe. Within the temporal parcellations, we found that *Te* occurred earliest in the superior temporal gyrus, followed by the medial and inferior temporal gyrus and lateral occipitotemporal gyrus. Interestingly, all the *Te* in the posterior parts of the temporal lobe occured earlier than the anterior part (Fig. 8a). Supplementary Table S1 shows the fitted model parameters in each cortical ROI. There were minor regional variations in peak cortical thickness *Ym*, with higher values in parietal lobe and temporal lobe and lower values in frontal lobe and occipital lobe. Within the temporal parcellations, *Ym* of the superior temporal gyrus was higher than those in medial and inferior temporal gyrus and lateral occipitotemporal gyrus.

**Fig. 4.**
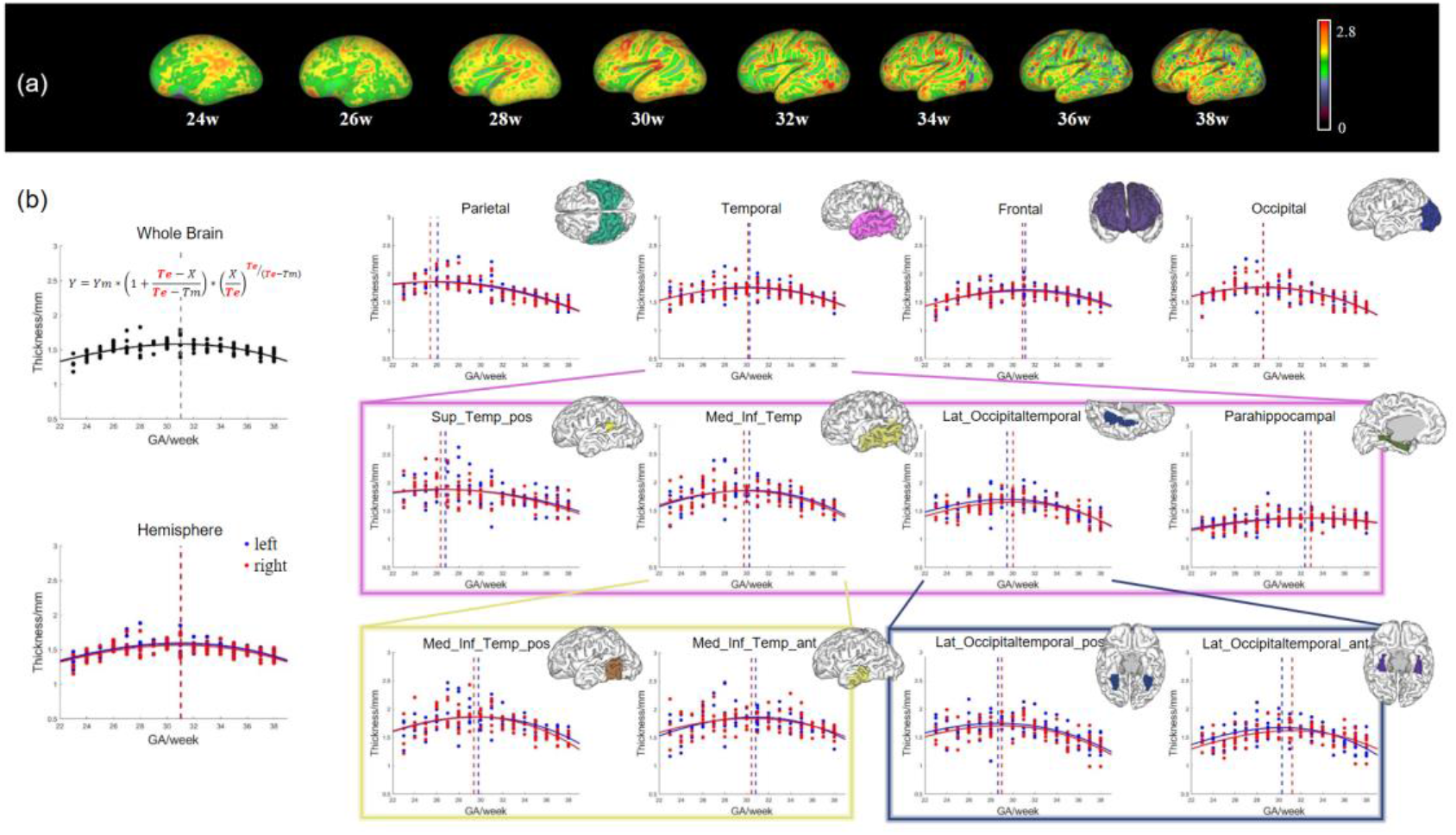
Developmental trajectories of the cortical thickness in the fetal brains. (a) The thickness on the inflated surface of atlas at every other GA week. (b) Beta growth and decay model fitting of cortical thickness averaged over the whole brain, hemispheres, and individual cortical regions. The dash line represents *Te* in different regions for the left (blue) and right (red) sides. The temporal lobe was further divided into four subregions in the pink box, including the superior temporal gyrus (only posterior part presented), medial inferior temporal gyrus, lateral occipitotemporal gyrus, and parahippocampal gyrus. The medial inferior temporal gyrus was further parcellated into the posterior part and anterior part (yellow box) and the lateral occipitotemporal gyrus was further divided into the posterior part and anterior part (blue box). Abbreviations: Sup=Superior, Temp=Temporal, Med=Medial, Inf=Inferior, Lat=Lateral, pos=posterior, ant=anterior.

**Fig. 5.**
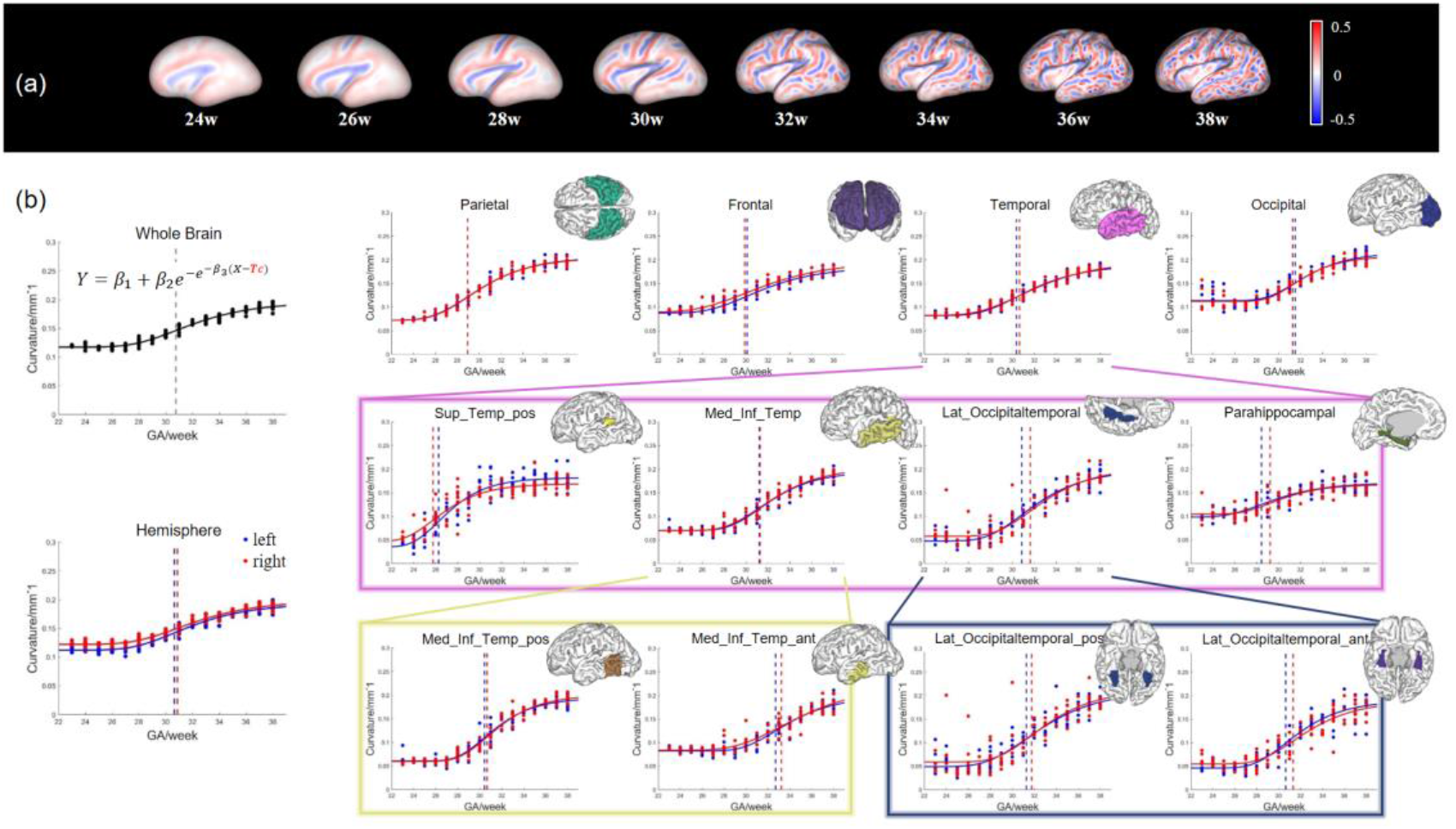
Developmental trajectories of the curvature in the fetal brains. (a) The curvature on the inflated surface of atlas at every other GA week. (b) Gompertz model fitting of curvature averaged over the whole brain, hemispheres, and individual cortical regions. The dash line represents *Tc* in different regions for the left (blue) and right (red) sides. The temporal lobe was further divided into four subregions in the pink box, including the superior temporal gyrus (only posterior part presented), medial inferior temporal gyrus, lateral occipitotemporal gyrus, and parahippocampal gyrus. The medial inferior temporal gyrus was further parcellated into the posterior part and anterior part (yellow box) and the lateral occipitotemporal gyrus was further divided into the posterior part and anterior part (blue box). Abbreviations: Sup=Superior, Temp=Temporal, Med=Medial, Inf=Inferior, Lat=Lateral, pos=posterior, ant=anterior.

**Fig. 6.**
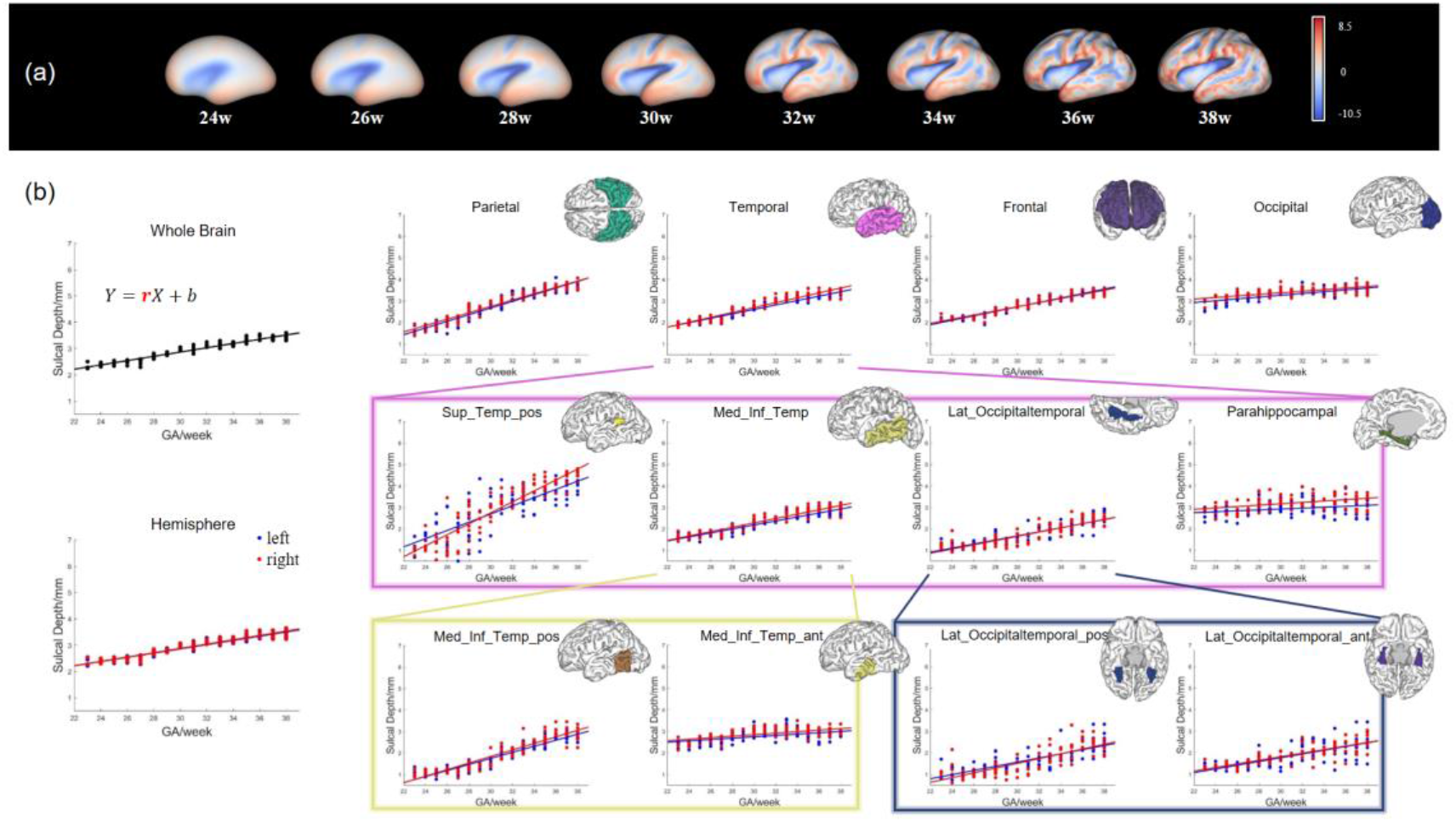
Developmental trajectories of the sulcal depth in the fetal brains. (a) The sulcal depth on the inflated surface of atlas at every other GA week. (b) Linear fitting of sulcal depth averaged over the whole brain, hemispheres, and individual cortical regions (blue – left hemisphere, red – right hemisphere). The temporal lobe was further divided into four subregions in the pink box, including the superior temporal gyrus (only posterior part presented), medial inferior temporal gyrus, lateral occipitotemporal gyrus, and parahippocampal gyrus. The medial inferior temporal gyrus was further parcellated into the posterior part and anterior part (yellow box) and the lateral occipitotemporal gyrus was further divided into the posterior part and anterior part (blue box). Abbreviations: Sup=Superior, Temp=Temporal, Med=Medial, Inf=Inferior, Lat=Lateral, pos=posterior, ant=anterior.

**Fig. 7.**
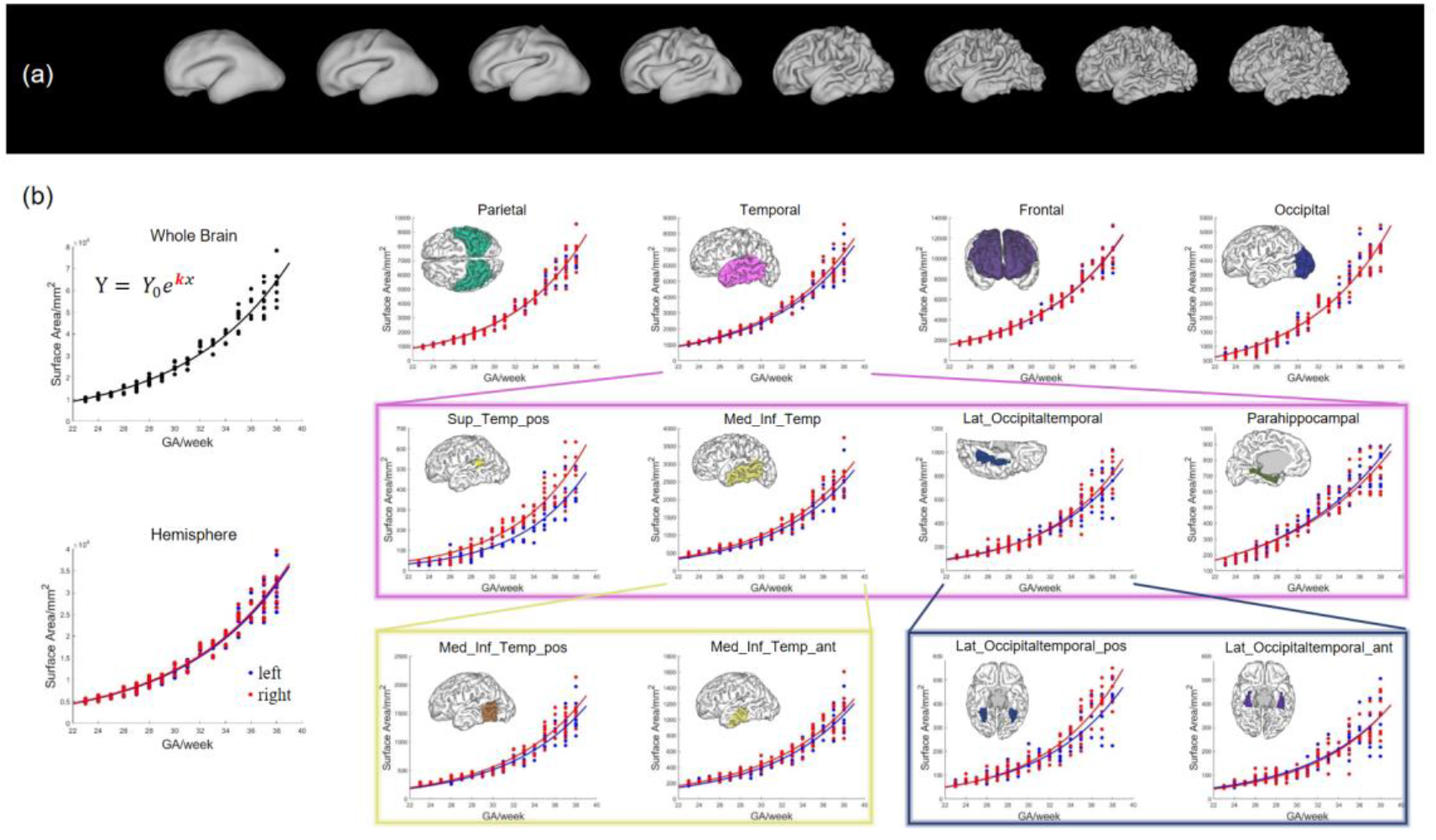
Developmental trajectories of the surface area in the fetal brains. (a) The surface area on the inflated surface of atlas at every other GA week. (b) Exponential growth model fitting of surface area averaged over the whole brain, hemispheres, and individual cortical regions (blue – left hemisphere, red – right hemisphere). The temporal lobe was further divided into four subregions in the pink box, including the superior temporal gyrus (only posterior part presented), medial inferior temporal gyrus, lateral occipitotemporal gyrus, and parahippocampal gyrus. The medial inferior temporal gyrus was further parcellated into the posterior part and anterior part (yellow box) and the lateral occipitotemporal gyrus was further divided into the posterior part and anterior part (blue box). Abbreviations: Sup=Superior, Temp=Temporal, Med=Medial, Inf=Inferior, Lat=Lateral, pos=posterior, ant=anterior.

**Fig. 8.**
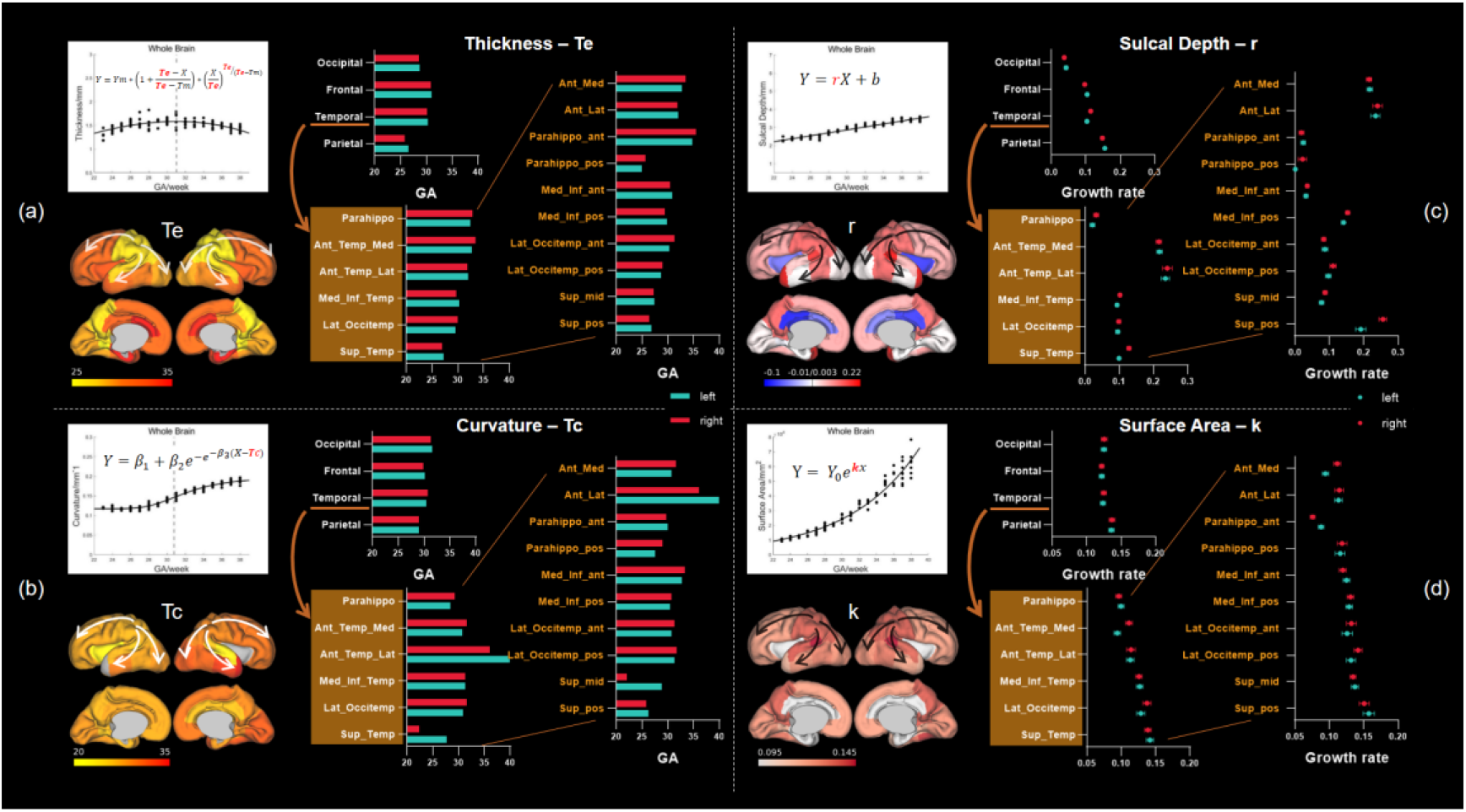
Spatiotemporal pattern of cortical development based on the critical change time of the cortical thickness (*Te* in a) and curvature (*Tc* in b) and the growth rate of the sulcal depth (*r* in c) and surface area (*k* in d). Scatter plots in the upper left corner of (a-d) show the model fitting results of the whole-brain averaged cortical morphological indicators. The corresponding parameter maps in the bottom left (*Te* of the Beta growth and decay model, *Tc* of the Gompertz model, *r* of the linear model, and *k* of the exponential model) are shown in the medial and lateral views of midthickness surface of a 30 week atlas. The bar plots or dot plots in the right panels of (a-d) compare the model parameters among parietal, temporal, frontal, and occipital lobes and different subregions within the temporal lobe. Abbreviations: Sup=Superior, Temp=Temporal, Med=Medial, Inf=Inferior, Lat=Lateral, pos=posterior, ant=anterior, Occipitemp=Occipitotemporal, Parahippo=parahipppocampal.

#### Nonlinear development of cortical curvature

Curvature showed a nonlinearly increasing trend across GA (Fig. 5) that was consistent with the Gompertz model. The *Tc* in the Gompertz model indicated the GA where the peak growth rate occurred, and *Tc* of the whole brain was around 31 weeks GA, similar to the turning point in cortical thickness. Fig. 5b shows the development order of *Tc* in different regions, which was similar to that of cortical thickness, following the order of parietal lobe, frontal lobe, temporal lobe, and occipital lobe. The *Tc* order among temporal lobe parcellations and anterior-posterior gradient (Fig. 8b) was consistent with *Te* of cortical thickness. Supplementary Table S2 shows the fitted Gompertz model parameters in each cortical ROI.

#### Linear increase of cortical gyrification

Sulcal depth that indicates the degree of gyrification showed a linear increase over observed GA (Fig. 6). The slope *r* in the linear model indicates the growth rate of sulcal depth. Fig. 6b shows various growth rates in different regions, with faster rates in parietal and temporal lobes than those in the frontal and occipital lobes. This order of growth rate matched with the order of *Tc* time in curvature analysis. The fastest increase of sulcal depth observed in parietal lobe corresponded to early development of the primary and secondary sulcus such as central sulcus and postcentral sulcus. The occipital lobe had the deepest sulcus at early GA but fell behind later due to the slow growth. Among the temporal parcellations, we also found that the sulcal depth of the posterior part increased faster than the anterior part (Fig. 8c). The posterior part of the superior temporal gyrus which is also known as Wernicke’s area grew rapidly and showed obvious developmental asymmetry during the observed GA period, whereas parahippocampal gyrus showed a relatively stable trajectory.

#### Exponential expansion of surface area

Fig. 7 shows that cortical surface area followed an exponentially increasing trend across GA, which is consistent with previous studies^12,15^. The *k* parameter in exponential growth model indicates the growth rate of surface area. Supplementary Table S4 shows that parietal lobe expanded fastest, followed by the occipital lobe, temporal lobe, and frontal lobes, which agreed with *Te* order of cortical thickness. The order of growth rate in the temporal parcellations and the anterior-posterior gradient (Fig. 8d) were also the same as patterns of *Te* in cortical thickness and *Tc* in curvature. In addition, we observed substantial asymmetry in several temporal regions compared to the other cortical indictors.

Overall, the results from cortical thickness, curvature, gyrification, and surface area collectively pointed to a central-to-peripheral developmental gradient that parietal lobe developed the earliest and fastest, followed by temporal, frontal, and occipital lobes. Within the temporal lobe, the superior temporal gyrus developed the earliest and fastest, followed by the medial and inferior temporal gyrus and lateral occipitotemporal gyrus. Beyond the superior-to-inferior gradient, we also found a posterior-to-anterior pattern in the development of temporal lobe. Particularly, the cortical thickness and surface area shared the same developmental order. These spatiotemporal patterns of fetal cortical development were summarized in Fig. 8.

### 3.3 Structural asymmetries

The developmental trajectories in section 3.2 indicated laterality in many regions of the fetal brain. Some structural asymmetries in gyri and sulci can be visually appreciated on T2w brain images (arrows and circles in Fig. S3). Quantitative analysis revealed significant asymmetric regions in terms of surface area (Fig. 9), curvature (Supplementary Fig. S4), and sulcal depth (Supplementary Fig. S5) as indicated by yellow outlines.

**Fig. 9.**
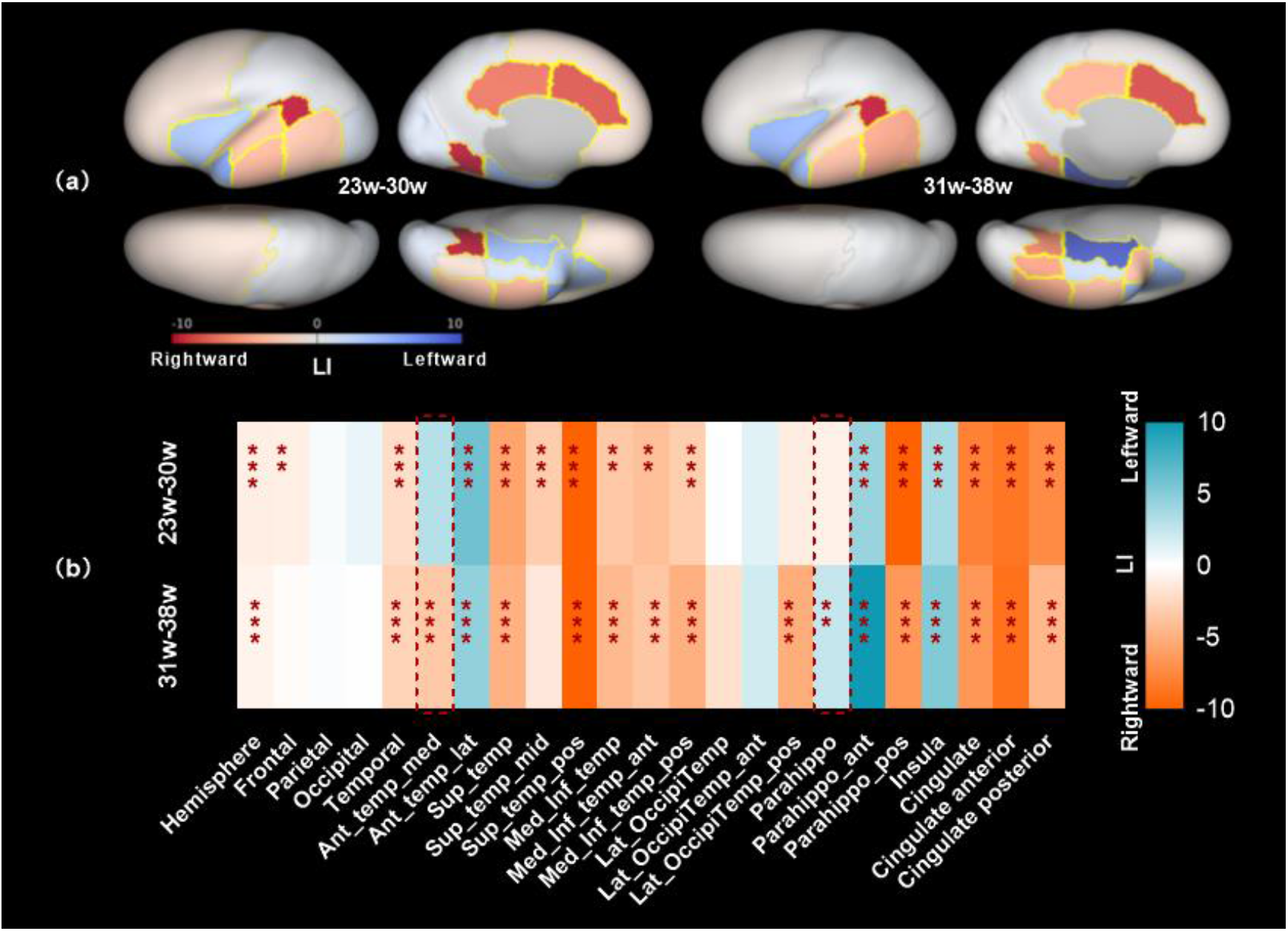
Cortical asymmetry in terms of surface area. (a) LI maps of the fetal brain in the 23-30w and 31-38w groups. Positive LI indicates leftward asymmetry and negative LI indicates rightward asymmetry. Significantly asymmetric regions are indicated by yellow outlines. (b) Statistical analysis of the LIs in the two GA groups. Red dashed box indicated the regions in which lateralization reversed between the two GA groups. *Adjusted P <0.05, **adjusted P<0.01, ***adjusted P <0.001. Abbreviations: Sup=Superior, Temp=Temporal, Med=Medial, Inf=Inferior, Lat=Lateral, pos=posterior, ant=anterior, Occipitemp=Occipitotemporal, Parahippo=parahipppocampal.

Regarding surface area, the frontal, parietal, and occipital lobes did not exhibit strong asymmetry, whereas many subregions in the temporal lobe did. Two language-related regions, the posterior part of superior temporal gyrus (STG, aka. Wernicke’s area) and the anterior part of STG, presented prominent rightward asymmetry with negative LI during the fetal period. But another language-related region, the insula, demonstrated leftward asymmetry. Within the parahippocampal gyrus that is related to visuospatial processing, we found leftward asymmetry in the anterior part, whereas rightward asymmetry was observed in the posterior part. This pattern can also be found in asymmetry of curvature (Supplementary Fig. S4). Moreover, the visual-temporal region and medial and inferior temporal gyrus demonstrated significant rightward asymmetry, which was also found in sulcal depth (Supplementary Fig. S5). Overall, there was a general leftward lateralization in the posterior region and a rightward asymmetry in the anterior brain in terms of the surface area and curvature.

Along the timeline, the 23-30w and 31-38w groups exhibited similar asymmetry pattern, expect for some of the regions that shifted laterality with GA (red dashed box in Fig. 9b and Supplementary Figs. S4-S5).Curvature of the posterior part of the STG presented negative LI (rightward) in the early period but changed to positive LI (leftward) in the late period (Supplementary Fig. S4), while the sulcal depth in this region showed the opposite trend (leftward to rightward asymmetry, Supplementary Fig. S5).

### 3.4 Cortical morphological differences between CHN and CRL atlases

Visual comparison between CRL atlas^19^ and our CHN atlas indicated regional differences in a few anatomical details, e.g., more convoluted gyrification in the temporal and occipital lobes of the CHN atlas compared with CRL (red circles and arrows in Fig. 10a and Supplementary Fig. S6), and their developmental trajectories (Fig. 10c) supported this observation. For instance, the CRL atlas showed slightly higher cortical thickness but lower curvature than the CHN brains between 27-35w of GA. Fig. 10d and Supplementary Fig. S7a show the morphological differences between our atlas and CRL atlas using TBM-based log-Jacobian determinant, which showed the most prominent difference in the central sulcus and parietal lobe in early period and also occipital lobe and cerebellum at later GA (arrows in Fig. 10d and Fig. S7a). Furthermore, the TBM-based growth vector fields obtained by transforming the CHN atlas to CRL atlas at corresponding GAs (Fig. 10b and Supplementary Fig. S7b) indicated the gyri of CHN brains were more convex and the sulci were deeper, e.g., the growth vectors on gyri pointed outward and the growth vectors on sulci pointed inward. Note that statistical comparison was not performed since we do not have the individual fetal brains from CRL.

**Fig. 10.**
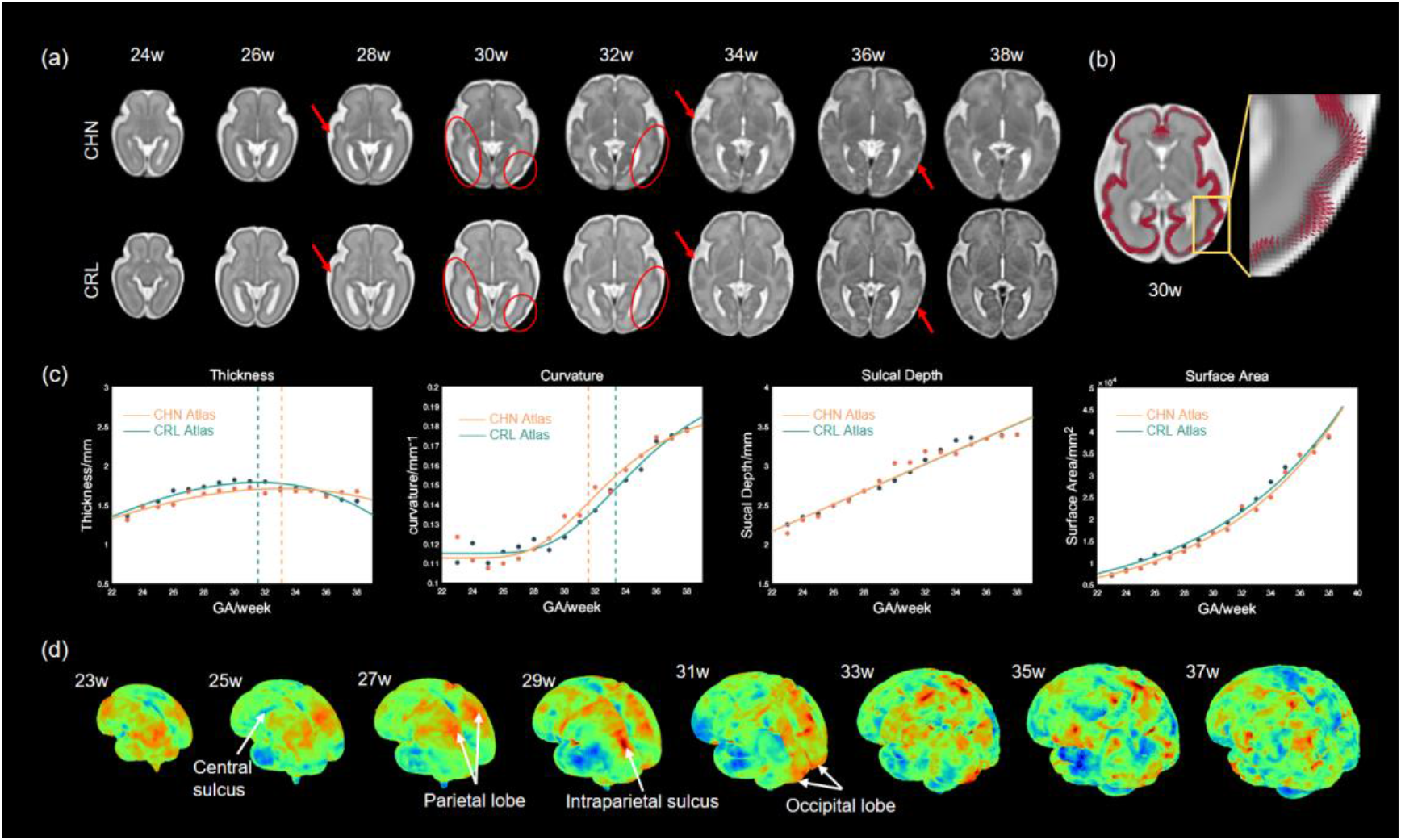
Comparison between CHN atlas and CRL atlas. (a) Visual comparison of T2w images at matched sections between CHN atlas and CRL atlas at representative GAs. The primary differences were indicated by red circles and arrows showing that our CHN atlas had more convoluted gyrification in temporal lobe. (b) The deformation fields obtained by transforming CHN atlas to CRL atlas at 30w GA. (c) Trajectories of thickness, curvature, sulcal depth, and surface area over GA of our CHN atlas (orange) and CRL atlas (green). (d) Morphological differences between CHN atlas and CRL atlas based on the logarithm of the Jacobian determinant from SyN registration between CHN and CRL atlases. The colors in red and yellow represent morphological expansion, indicating more convex gyrus in CHN atlas than CRL atlas, and green and blue show morphological shrinkage, indicating deeper sulcus in CHN atlas the CRL atlas.

## 4 Discussion

In this study, we generated a high-definition spatiotemporal structural MRI atlas of the fetal brain based on a Chinese population, aiming to preserve anatomical details in the population-average templates using adaptive kernel regression, pairwise initialization, shape update and iterative SyN registrations. Moreover, this study systematically quantified the spatiotemporal growth pattern of cortical features, including the biphasic changes in cortical thickness, nonlinear increase in curvature, linear growth of sulcal depth, and exponential expansion of surface area. Based on the characteristics of these trajectories, the developmental order of different cortical regions was mapped, which collectively pointed to a unique parietal-to-peripheric gradient. We also characterized the inter-hemisphere asymmetry began in early fetal period and it developed with GA in terms of degree and laterality. These findings provided a comprehensive view of normal early human brain development in utero.

### 4.1 The Chinese fetal brain atlas

High-quality fetal brain atlases based on Caucasian or mixed populations have been developed by Gholipour et al.^19^, Serag et al.^18^, etc. Previous studies have found considerable morphological and microstructure differences in human brains between different populations due to genetic and environmental disparities, both in the adult and pediatric brains^22,41^. Wu et al. found the anatomical differences in several cortical regions between their proposed Chinese fetal brain atlas compared to the Caucasian population atlas^12^. In addition, several studies have demonstrated the necessity to build a population-specific based brain atlas^42–44^ for quantitative analysis, such as segmentation. However, the existing Chinese fetal brain atlases^12,24^ are still insufficient, which both utilized clinical quality data at 1.5T with thick slices (4-5mm), making it difficult to be directly compared with the other atlases. The proposed Chinese fetal brain atlas was built on research protocol data acquired at 2mm thick-slices, multiple 2D stacks (3-9 stacks per subject), and a wide GA range from 23w to 38w. We used an adaptive kernel algorithm to compute the average brain template using weighted images from the temporal neighbors of the target age^30^. This technique also equalizes the distribution of individuals across selected ages. Besides, we used shape update based pairwise registration and iterative groupwise registration method to find the minimum shape distance in diffeomorphic space which determines the optimal unbiased shape^45^.

This study identified several morphological differences between the CHN and CRL fetal brain atlases, e.g. deeper gyrification but thinner cortical thickness in our CHN atlas compared to CRL atlas, indicating potential ethnic difference in the fetal brain, possibly related to genetic factors before environmental factors come in. It is worth noting that in addition to biological effect, the difference in image contrast from different acquisition protocols can also contribute to the observation. Further studies require individual data from both atlases with the data harmonization and consistent reconstruction pipelines for a fair statistical comparison.

### 4.2 Morphological development of the fetal brains

#### Biphasic change of cortical thickness and developmental order

The changes in cortical morphological properties reflect the underlying cellular events during development. In this study, we found an interesting biphasic development pattern of cortical thickness across the observed fetal period. The same trend was replicated in the CRL atlas which was processed with dHCP-structural-pipeline based on CRL segmentation (Fig. 10c). To the best of our knowledge, developmental trajectories of cortical thickness in infants, adolescents, and adults have been well delineated^46–51^, but its development in fetal period was rarely characterized. Xu et al. depicted the cortical thickness trajectory of 14w to 22w GA postmortem fetuses, showing a trend of gradual increase^9^. The age range of this study may be integrated with our study to complete the timeline. We think the ascending curve corresponds to the stage when a large number of neuronal cells migrate from proliferate zone to the cortical plate and proliferate in the cortical plate^52^. It has been suggested that neuronal migration and proliferation are vigorous in the second trimester, and the process gradually ceases during the third trimester^1,2,52^.

The ascending trend was terminated by a turning point that varies in different brain regions, around 30-32w of GA. The turning point coincided with the timing of dendritic arborization of the post-migratory neurons in the cortical plate as well as the thalamocortical fibers afferent infiltration from subplate to cortical plate^2,52–55^. Diffusion MRI studies^56,57^ indicated that the cortical ingrowth of thalamocortical afferent fibers and tangential increase in dendritic arborization are two major causes of diffusivity decrease that takes place around 30-31 weeks GA which matched with our findings. Zhang et al.^58^ demonstrated in postmortem fetal brains that there was a high signal layer in T1w image during 24-26 weeks where the thalamocortical fibers reside. This layer gradually became indistinguishable with cortical plate as the thalamocortal fibers migrated to the cortical plate around 28 weeks, and thereafter, the cortical plate appeared thinner as the further ingrowth of the thalamocortical fibers^58^. Although the in-utero MRI at 3T scanner had relatively low resolution compared to the postmortem study, it is possible that the images captured this process as a change in image contrast. The turning point in cortical thickness occurred the earliest in the parietal lobe where the somatosensory cortex locates, followed by the occipital, temporal, and frontal lobes in our results. The observation agreed with the fact that thalamocortical afferent in somatosensory develops 1-2 weeks earlier than occipital and frontal lobes^59^.

#### Radial growth versus tangential expansion of cortical plate

After neurons migrate to their destination in the cortical plate, there is a gradual acceleration of dendritic differentiation in the cortical plate during the third trimester^1,52,55^. The vast expansion of dendritic arborization in the tangential direction of the cortical plate is associated with the surface area expansion^40,52^. This probably explains our results that the order of growth rates in surface area matched with the turning points of cortical thickness across different cortical lobes and subregions. Vasung et al. demonstrated that the cortical surface increased by almost 50 times from 15 to 42 weeks GA, contrasting a subtle sub-millimeter increase in its thickness^52^. Tallinen et al. reported that the 2-fold increase in thickness is very small compared with the 30-fold increase in surface area^60^. Therefore, the tangential growth of dendritic arborization could be much more drastic compared to the radial growth in the cortical plate, resulting in a relative decline in cortical thickness at later gestation.

As summarized in Fig. 11, we hypothesize that the major cellular events correlated with the development of cortical thickness are neuronal migration and proliferation, thalamocortical fibers afferent, and dendritic differentiation. The mixed contributions from these processes could explain the spatiotemporal changes of various cortical morphological indicators in this study. We are aware that the correlation of these events with MRI findings is only based on indirect inferences, and therefore further direct evidence is needed to confirm this hypothesis.

**Fig. 11.**
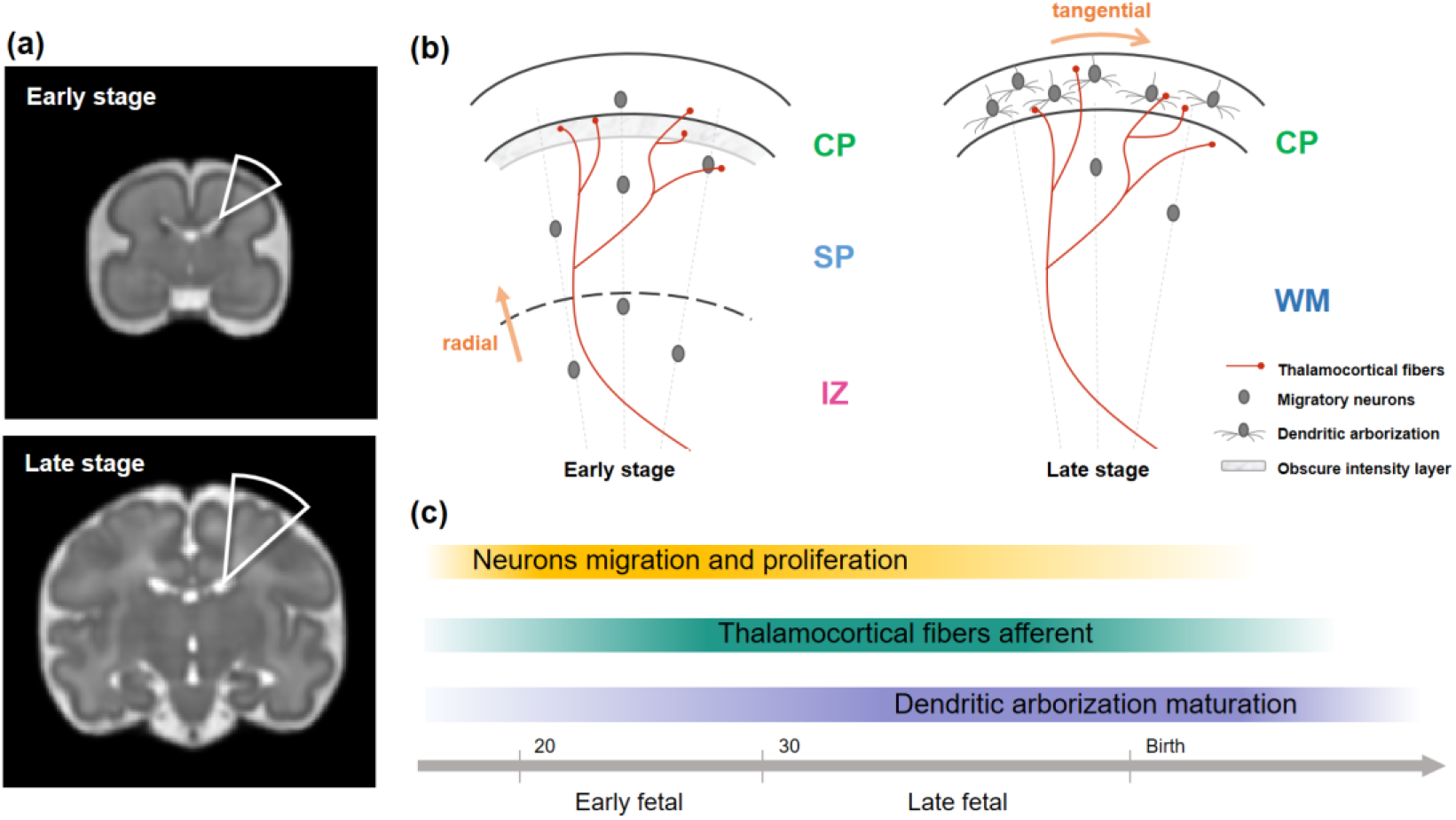
Schematics of the key neurological cellular events related to the developmental changes of cortical thickness in fetuses. (a) Representative fetal brain templates at an early stage (24w) and late stage (35 weeks). (b) Schematics of the hypothesized processes of neuronal migration and proliferation, thalamocortical fibers ingrowth, and dendritic arborization differentiation that are related to our MRI findings during the early and late periods. (c) Intensity of these processes from prenatal to postnatal stages summarized from the current and previous studies. Abbreviations: CP=cortical plate, SP=subplate; IZ=intermediate zone; WM=white matter.

#### Development of gyrification and its driving forces

Previous studies have demonstrated that cortical folding increases intensely in the third trimester, following a specific timetable in different regions^8,14,61,62^. Here, we computed the curvature, sulcal depth, and surface area to investigate the cortical gyrification from 23w to 38w GA. The developmental trend of curvature in our results demonstrated a slow increase initially, followed by a period of rapid growth that slowed down again towards birth. This type of growth trend can be best modeled by the Gompertz function^8^. We found that the growth rate of sulcal depth and the peak growth time point of curvature pointed to the same development order from parietal lobe to temporal, frontal, and occipital lobes. Also using the Gompertz model, Wright et al. showed that gyrification accelerates rapidly between 25 and 30 weeks and that posterior temporal lobe and parietal lobe had the highest peak growth rate while the frontal lobe developed slower^8^, which is consistent with our results. The occipital lobe showed the slowest growth rate in sulcal depth and latest peak growth rate in curvature but it had the higher sulcal depth and curvature at the beginning (23-24w), possibly because the occipital lobe had fewer main gyri and sulci compared with other lobes, except for the early developed Parieto-occipital sulcus and Calcarine sulcus around 23-24w.

There are two dominant theories for the genesis of cerebral cortex folding. Richman et al.^63^ put forth the “buckling due to differential expansion” theory, which stated that compressive forces arise in the outer layer as a consequence of its tangential expansion relative to the inner zone and these forces lead to buckling of the cortex. Van Essen^64^ proposed the “axon tension” theory in which posited tension from axons in the inner zone drives cortical folding by pulling gyral walls toward one another. Based on the good agreement between the turning time point of cortical thickness and the timing of maximum growth rate in curvature as well as the growth rate order of surface area in different regions, we think that the cortical thickness decrease after the turning point and surface area expansion were related to dendritic differentiation in tangential direction, and thus supporting the “buckling theory”. Moreover, the regional variability of cortical folding observed in our results require further investigation. Lui et al.^65^ and Rellio et al.^66^ indicated that nonuniform distribution of radial glial progenitor cells that support the tangential dispersion may cause the region specification in cortical folding, while Vasung et al. suggested that the regional specification can be described by protomap (regional destiny of cortical neurons and the size of cortical area are genetically determined) and protocortex (regional growth rate are different and are shaped by external influence) hypotheses^16^. Computational or mechanical models^60,67,68^ that connect these biological processes and folding patterns may help to support these theories.

### 4.3 Developmental asymmetry

Asymmetry in cell size, columnar organization, and the complexity of dendritic arbors hidden under morphological indicators have been reported^69,70^, and these microscopic biological processes are reflected in the asymmetry of cortical morphology. Brain morphological asymmetries in adults and infants have been well portrayed in previous studies^71–77^. How early brain asymmetry starts and how it changes during fetal stage remains unclear. There are a few in-vivo fetal studies that showed hemispheric differences in the volume of different fetal brain regions^5–7,78^. Habas et al.^4^ showed the asymmetry of cortical curvature in functionally specialized areas between 20-28w GA. Kasprian et al.^79^ analyzed the size and shape asymmetry of temporal lobe and some major sulci in temporal lobe. Here we presented a comprehensive picture of the cortical asymmetry in healthy fetal brains. Specially, we found that STG demonstrated rightward asymmetry in surface area which has also been reported in fetuses^4,79^, neonates^72,80^, and infants^81^, while Kong et al. found leftward asymmetry of the STG in adults^71^. Our results showed that the lateralization index of the middle part of STG decreased with development, suggesting that there may be a developmental shift of asymmetry in the STG. In contrast, the insula, also a language related area, demonstrated leftward asymmetry in surface area, which already shared adult brain pattern^71^. It is well-known that one of the most striking features in human brains is an overall leftward asymmetry in the occipital lobe and a rightward asymmetry in frontal lobe (or brain torque)^82^, which is also seen in our data. The lateralization index of surface area in the visual related region of occipital lobe was positive, indicating a leftward tendency, although there was no statistical significance.

Interestingly, several regions were found to exhibit asymmetry patterns similar to that in neonates^72,76,81,83^. For example, the medial part of anterior temporal lobe which related to emotion demonstrated the rightward lateralization in late period in surface area that agreed with the neonates study^72^. The parahippocampal gyrus exhibited regional variability with leftward asymmetry in the anterior part and rightward asymmetry in the posterior part in surface area. This phenomenon was also observed in a neonatal study^72^, possibly suggesting that the subregions of the parahippocampal gyrus may be involved in different functions. An adult functional MRI study showed that the posterior section of the parahippocampal cortex was predominantly associated with pictorial scene analysis while the anterior section was related to contextual judgments^84^.

### 4.4 Limitation

In addition to the lack of direct evidence linking cortical developmental changes to biophysical events as mentioned above, there are some other limitations in this study. The number of T2w image data at large GA was still small for atlas construction, although we have used an adaptive kernel regression algorithm^12,18,30,49^ to compensate for the uneven data distribution. Besides, the DrawEM toolbox that we used to segment the fetal brain gave relatively large parcellations (except for the temporal lobe), which is not ideal for mapping the regional variability. Better fetal brain parcellation atlases are desired for detailed analysis. Moreover, the segmentation results from automated pipeline was not accurate enough and had to be manually refined, which is time-consuming and is subjective to manual errors. More advanced segmentation methods or deep learning approaches may help in this regard.

## 5 Conclusions

We generated a 4D spatiotemporal atlas of the normal fetal brain development from 23 to 38 gestational weeks in a Chinese population, based on which, we quantified the developmental gradient of fetal cortex. The parametrization of developmental trajectories indicated a critical period during fetal brain development around 31w GA, and also revealed a central-to-peripheral gradient of cortical development, which can be linked to biological cellular events during fetal development and possibly explained the cortical folding theory. Also, we found the majority of cortical regions already exhibited significant asymmetry during the early fetal period. The fetal brain atlas and well-characterized developmental patterns of fetal cortex may facilitate our understanding of fetal brain development in utero and can be potentially used for prenatal diagnostic purposes.

