## Supplementary for "Spatiotemporal atlas of the fetal brain depicts cortical developmental gradient in Chinese population"

### Supplementary Material

#### Supplementary Tables 1-5

**Table S1** Summary of existing fetal brain atlases up to date.

| Atlas | Atlas Information | | | Atlas Generation method | | | Number of participants | Acquisition details |
|---|---|---|---|---|---|---|---|---|
| | Modality | GA range | Resolu-tion | Registration | Atlas construction | Kernel regression | | |
| (Habas, Kim et al. 2010) | T2w | 21 to 24 weeks | 0.5mm isotropic | Elastic deformation based on tissue maps | Group-wise | No | 20 healthy fetuses | 1.5 T 1x1x3 $mm^3$ |
| **BioMedIA** (Serag, Aljabar et al. 2011) http://brain-development.org/ | T2w | 23 to 37 weeks | 1.18mm isotropic | Affine + Bspline free form deformation | Pairwise | Adaptive Gaussian kernel | 80 healthy fetuses | 1.5T 1.25×1.25×2.5$mm^3$ |
| (Zhan, Dinov et al. 2013) http://www.loni.ucla.edu/Atlases/ | Postmortem T2w (Chinese population) | 15 to 22 weeks | – | SyN in ANTs | Group-wise | No | 34 postmortem fetal brains | 7T 0.19×0.19×0.5$mm^3$ or 0.23×0.23×0.5$mm^3$ |
| (Dittrich, Riklin Raviv et al. 2014) Not publicly available | Ventricle only | 20 to 30 weeks | 0.78-0.9mm isotropic | Semi-supervised learning | – | Gaussian kernel | 32 healthy fetuses + 12 lissencephaly fetuses | 1.5T 0.78-0.9 mm in-plane, 3-4.4mm slice thickness |
| (Wright, Makropoulos et al. 2015) Not publicly available | Cortical Surface Atlas | 23 to 37 weeks | – | Surface registration based on sulcal alignment, | – | Gaussian kernel | 80 healthy fetuses | 1.5T |
| **CRL Atlas** (Gholipour, Rollins et al. 2017) http://crl.med.harvard.edu/research/fetal_brain_atlas/ | T2w | 21 to 37 weeks | 0.8mm isotropic | Greedy SyN in ANTs | Group-wise | Fixed-width Gaussian kernel | 81 healthy fetuses | 3T or 1.5T 0.9~1.1mm in-plane, 2mm slice thickness |
| (Xia, Wang et al. 2019) Not publicly available | Cortical Surface Atlas | 26.5, 27, 28.2, 28.7 weeks | | Surface registration | – | No | 25 normal fetuses | 1.5T 0.5×0.5×2.5 $mm^3$ |
| (Li, Yan et al. 2021) Not publicly available | T2w (Chinese population) | 23 to 36 in every 2 weeks | 0.8mm isotropic | Affine + SyN in ANTs | Pairwise + group-wise | No | 212 fetal brains collected, 35 for atlas | 1.5T 0.74×0.74×4$mm^3$ |
| **FBA atlas** (Wu, Sun et al. 2021) https://github.com/DeepBMI/ FBA-Chinese | T2w (Chinese population) | 22 to 35 weeks | 0.8mm isotropic | SyN in ANTs | Group-wise | Adaptive Gaussian kernel | 735 fetal brains collected, 115 for atlas | 1.5T 4mm slice thickness |
| **dHCP** (Uus, Matthew et al. 2021) To be deposited on https://gin.g-node.org/SVRTK | T2w (craniofacial reserved) | 21 to 36 weeks | 0.7mm isotropic | MIRTK atlas generation pipeline | – | Gaussian kernel | 291 normal fetal brains collected, 190 best quality data for atlas | 3T 1.1×1.1×2.2 $mm^3$ |
| **Proposed atlas** https://github.com/Thea-Eddie-Amy/CHN-fetal-brain-atlas | T2w (Chinese population) | 23 to 38 weeks | 0.8mm isotropic | ANTs Rigid+affine+SyN Shape update | Pairwise +group-wise | Adaptive Gaussian kernel | 219 healthy fetal brains collected, 90 best quality data for atlas | 3T 1.1×1.1×2 $mm^3$ |

Abbreviations: GA = gestational age; SyN = symmetric diffeomorphic deformable registration algorithm;

ANTs = advanced normalization tools.

**Table S2** The turning point time (Te) from Beta growth and decay model fitting of cortical thickness in different cortical regions, and the adjusted p-value and $R^2$ for the goodness-of-fit of Te.

| Structure | | Te (weeks) | P value | $R^2$ |
|---|---|---|---|---|
| Global | | 30.96 | <0.001* | 0.348 |
| Hemisphere | left | 30.98 | <0.001* | 0.340 |
| | right | 30.94 | <0.001* | 0.333 |
| Parietal lobe | left | 26.46 | <0.001* | 0.513 |
| | right | 25.77 | <0.001* | 0.549 |
| Frontal lobe | left | 31.05 | <0.001* | 0.303 |
| | right | 30.81 | <0.001* | 0.282 |
| Occipital lobe | left | 28.65 | <0.001* | 0.392 |
| | right | 28.59 | <0.001* | 0.445 |
| Temporal lobe | left | 30.27 | <0.001* | 0.304 |
| | right | 30.11 | <0.001* | 0.321 |
| Ant_temp_med | left | 32.73 | <0.001* | 0.213 |
| | right | 33.42 | <0.001* | 0.364 |
| Ant_temp_lat | left | 31.94 | <0.001* | 0.291 |
| | right | 31.92 | <0.001* | 0.196 |
| Sup_temp_pos | left | 25.65 | <0.001* | 0.195 |
| | right | 26.35 | 0.296 | 0.298 |
| Sup_temp_mid | left | 27.00 | <0.001* | 0.255 |
| | right | 24.97 | <0.001* | 0.253 |
| Med_inf_temp_pos | left | 29.81 | <0.001* | 0.267 |
| | right | 29.39 | <0.001* | 0.381 |
| Med_inf_temp_ant | left | 30.81 | <0.001* | 0.207 |
| | right | 30.43 | <0.001* | 0.154 |
| Lat_Occipitaltemporal_pos | left | 28.66 | <0.001* | 0.331 |
| | right | 29.00 | <0.001* | 0.389 |
| Lat_Occipitaltemporal_ant | left | 30.29 | <0.001* | 0.290 |
| | right | 31.22 | <0.001* | 0.224 |
| Parahippocampal gyrus | left | 32.39 | <0.001* | 0.107 |
| | right | 32.92 | <0.001* | 0.163 |

Abbreviations: ant=anterior, Temp=Temporal, Med=Medial, Lat=Lateral, Sup=Superior, pos=posterior, mid=middle, Inf=Inferior.

**Table S3** The timing of peak growth (Tc) and growth rate ($\beta_3$) from Gompertz model fitting of cortical curvature in different cortical regions, and the adjusted p-value and $R^2$ for Tc and $\beta_3$, repectively.

| Structure | | Tc (weeks) | P value of Tc | $\beta_3$ ($mm^{-1}$/week) | P value of k | $R^2$ |
|---|---|---|---|---|---|---|
| Global | | 30.76 | <0.001* | 0.31 | <0.001* | 0.961 |
| Hemisphere | left | 30.65 | <0.001* | 0.32 | <0.001* | 0.954 |
| | right | 30.88 | <0.001* | 0.30 | <0.001* | 0.950 |
| Parietal lobe | left | 28.93 | <0.001* | 0.31 | <0.001* | 0.969 |
| | right | 28.93 | <0.001* | 0.31 | <0.001* | 0.970 |
| Frontal lobe | left | 30.11 | <0.001* | 0.28 | <0.001* | 0.937 |
| | right | 29.89 | <0.001* | 0.23 | <0.001* | 0.930 |

| | | | | | | |
|---|---|---|---|---|---|---|
| Occipital lobe | left | 31.52 | <0.001* | 0.37 | <0.001* | 0.882 |
| | right | 31.29 | <0.001* | 0.42 | <0.001* | 0.910 |
| Temporal lobe | left | 30.39 | <0.001* | 0.30 | <0.001* | 0.959 |
| | right | 30.66 | <0.001* | 0.26 | <0.001* | 0.955 |
| Ant_temp_med | left | 30.77 | <0.001* | 0.28 | 0.082 | 0.464 |
| | right | 31.57 | <0.001* | 0.53 | 0.137 | 0.358 |
| Ant_temp_lat | left | 39.99 | <0.001* | 0.02 | <0.001* | 0.608 |
| | right | 36.06 | <0.001* | 0.13 | 0.205 | 0.702 |
| Sup_temp_pos | left | 26.27 | <0.001* | 0.42 | <0.001* | 0.845 |
| | right | 25.77 | <0.001* | 0.39 | <0.001* | 0.784 |
| Sup_temp_mid | left | 28.84 | <0.001* | 0.26 | <0.001* | 0.852 |
| | right | 21.19 | 0.260 | 0.12 | 0.262 | 0.842 |
| Med_inf_temp_pos | left | 30.43 | <0.001* | 0.40 | <0.001* | 0.947 |
| | right | 30.68 | <0.001* | 0.38 | <0.001* | 0.960 |
| Med_inf_temp_ant | left | 32.71 | <0.001* | 0.33 | <0.001* | 0.922 |
| | right | 33.23 | <0.001* | 0.23 | <0.001* | 0.878 |
| Lat_Occipitaltemporal_pos | left | 31.28 | <0.001* | 0.27 | <0.001* | 0.864 |
| | right | 31.76 | <0.001* | 0.31 | <0.001* | 0.741 |
| Lat_Occipitaltemporal_ant | left | 30.65 | <0.001* | 0.34 | <0.001* | 0.888 |
| | right | 31.33 | <0.001* | 0.32 | <0.001* | 0.853 |
| Parahippocampal gyrus | left | 28.44 | <0.001* | 0.32 | <0.001* | 0.750 |
| | right | 29.23 | <0.001* | 0.39 | <0.001* | 0.749 |

Abbreviations: ant=anterior, Temp=Temporal, Med=Medial, Lat=Lateral, Sup=Superior, pos=posterior, mid=middle, Inf=Inferior.

**Table S4** Growth rate (r) from linear fitting of sulcal depth in different cortical regions, and its adjusted p-value and $R^2$.

| **Structure** | | **r (mm/week)** | **P value** | **$R^2$** |
|---|---|---|---|---|
| Global | | 0.081 | <0.001* | 0.929 |
| Hemisphere | left | 0.080 | <0.001* | 0.917 |
| | right | 0.082 | <0.001* | 0.924 |
| Parietal lobe | left | 0.154 | <0.001* | 0.897 |
| | right | 0.148 | <0.001* | 0.918 |
| Frontal lobe | left | 0.103 | <0.001* | 0.924 |
| | right | 0.096 | <0.001* | 0.909 |
| Occipital lobe | left | 0.042 | <0.001* | 0.458 |
| | right | 0.036 | <0.001* | 0.477 |
| Temporal lobe | left | 0.103 | <0.001* | 0.907 |
| | right | 0.113 | <0.001* | 0.897 |
| Ant_temp_med | left | 0.217 | <0.001* | 0.886 |
| | right | 0.215 | <0.001* | 0.887 |
| Ant_temp_lat | left | 0.234 | <0.001* | 0.800 |
| | right | 0.240 | <0.001* | 0.750 |
| Sup_temp_pos | left | 0.191 | <0.001* | 0.631 |
| | right | 0.255 | <0.001* | 0.836 |
| Sup_temp_mid | left | 0.077 | <0.001* | 0.575 |
| | right | 0.087 | <0.001* | 0.659 |
| Med_inf_temp_pos | left | 0.141 | <0.001* | 0.891 |
| | right | 0.153 | <0.001* | 0.860 |
| Med_inf_temp_ant | left | 0.031 | <0.001* | 0.216 |

| | | | | |
|---|---|---|---|---|
| | right | 0.035 | <0.001* | 0.372 |
| Lat_Occipitaltemporal_pos | left | 0.096 | <0.001* | 0.615 |
| | right | 0.110 | <0.001* | 0.625 |
| Lat_Occipitaltemporal_ant | left | 0.086 | <0.001* | 0.515 |
| | right | 0.083 | <0.001* | 0.658 |
| Parahippocampal gyrus | left | 0.021 | 0.003 | 0.077 |
| | right | 0.032 | <0.001* | 0.166 |

Abbreviations: ant=anterior, Temp=Temporal, Med=Medial, Lat=Lateral, Sup=Superior, pos=posterior, mid=middle, Inf=Inferior.

**Table S5** The fitted exponent (k) from Exponential growth model of surface area from different cortical regions, and its adjusted p-value and $R^2$.

| Structure | | k | P value | $R^2$ |
|---|---|---|---|---|
| Global | | 0.122 | <0.001* | 0.948 |
| Hemisphere | left | 0.122 | <0.001* | 0.947 |
| | right | 0.122 | <0.001* | 0.948 |
| Parietal lobe | left | 0.136 | <0.001* | 0.948 |
| | right | 0.137 | <0.001* | 0.941 |
| Frontal lobe | left | 0.122 | <0.001* | 0.943 |
| | right | 0.122 | <0.001* | 0.953 |
| Occipital lobe | left | 0.125 | <0.001* | 0.935 |
| | right | 0.125 | <0.001* | 0.930 |
| Temporal lobe | left | 0.123 | <0.001* | 0.941 |
| | right | 0.125 | <0.001* | 0.939 |
| Ant_temp_med | left | 0.094 | <0.001* | 0.838 |
| | right | 0.111 | <0.001* | 0.879 |
| Ant_temp_lat | left | 0.113 | <0.001* | 0.851 |
| | right | 0.114 | <0.001* | 0.821 |
| Sup_temp_pos | left | 0.157 | <0.001* | 0.864 |
| | right | 0.150 | <0.001* | 0.882 |
| Sup_temp_mid | left | 0.137 | <0.001* | 0.917 |
| | right | 0.134 | <0.001* | 0.943 |
| Med_inf_temp_pos | left | 0.128 | <0.001* | 0.907 |
| | right | 0.130 | <0.001* | 0.912 |
| Med_inf_temp_ant | left | 0.125 | <0.001* | 0.906 |
| | right | 0.112 | <0.001* | 0.888 |
| Lat_Occipitaltemporal_pos | left | 0.131 | <0.001* | 0.855 |
| | right | 0.141 | <0.001* | 0.902 |
| Lat_Occipitaltemporal_ant | left | 0.126 | <0.001* | 0.790 |
| | right | 0.132 | <0.001* | 0.830 |
| Parahippocampal gyrus | left | 0.099 | <0.001* | 0.885 |
| | right | 0.096 | <0.001* | 0.871 |
| Insula | left | 0.090 | <0.001* | 0.898 |
| | right | 0.095 | <0.001* | 0.872 |
| Cingulate | left | 0.076 | <0.001* | 0.760 |
| | right | 0.069 | <0.001* | 0.752 |

Abbreviations: ant=anterior, Temp=Temporal, Med=Medial, Lat=Lateral, Sup=Superior, pos=posterior, mid=middle, Inf=Inferior.

#### Supplementary figures S1-7

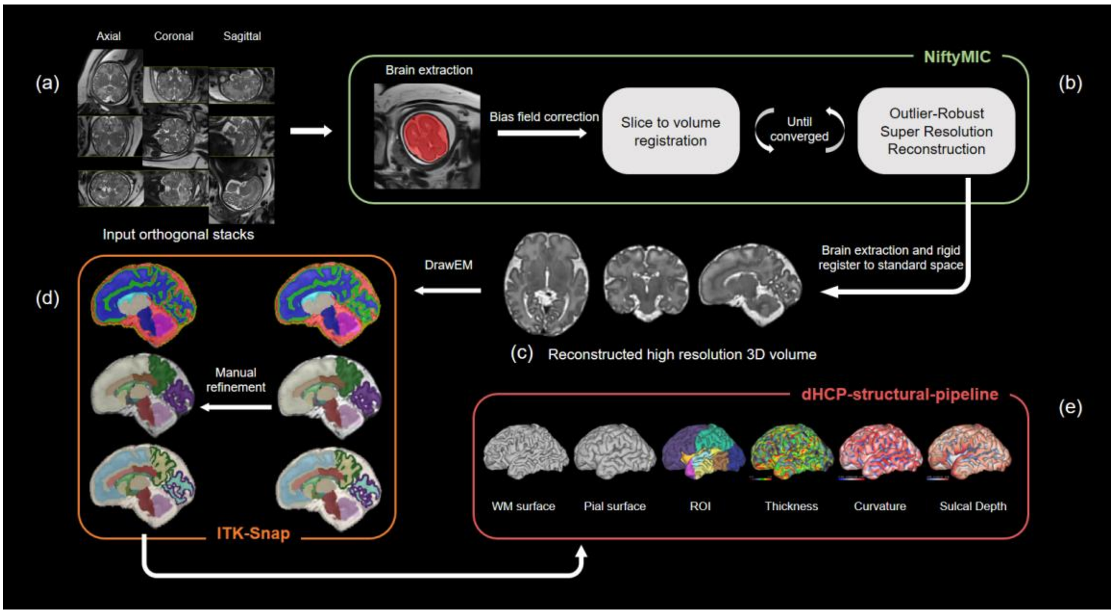


Fig. S1. Overview of fetal brain MRI data processing pipeline: (a) shows the original T2w HASTE scan in axial, coronal, and sagittal planes; (b) shows the major steps of NiftyMIC reconstruction pipeline; (c) is the reconstructed high-resolution 3D volume; (d) shows the segmentation results; (e) is the cortical analysis results in surface space.

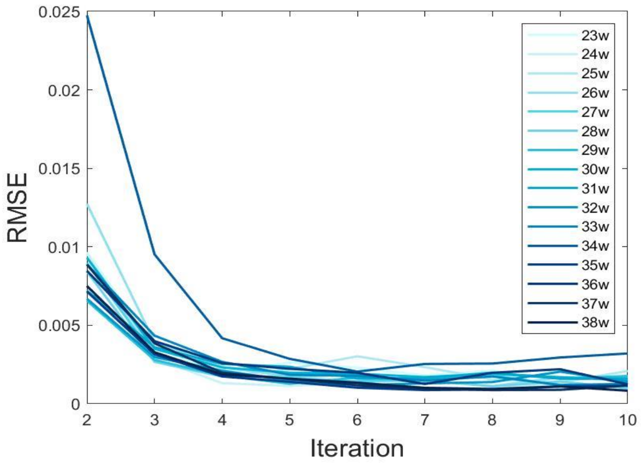


Fig. S2. Convergence of the iterative regrestration processes during atlas generation at different GA, represented by the root mean square error (RMSE) between the previous and current iteration.

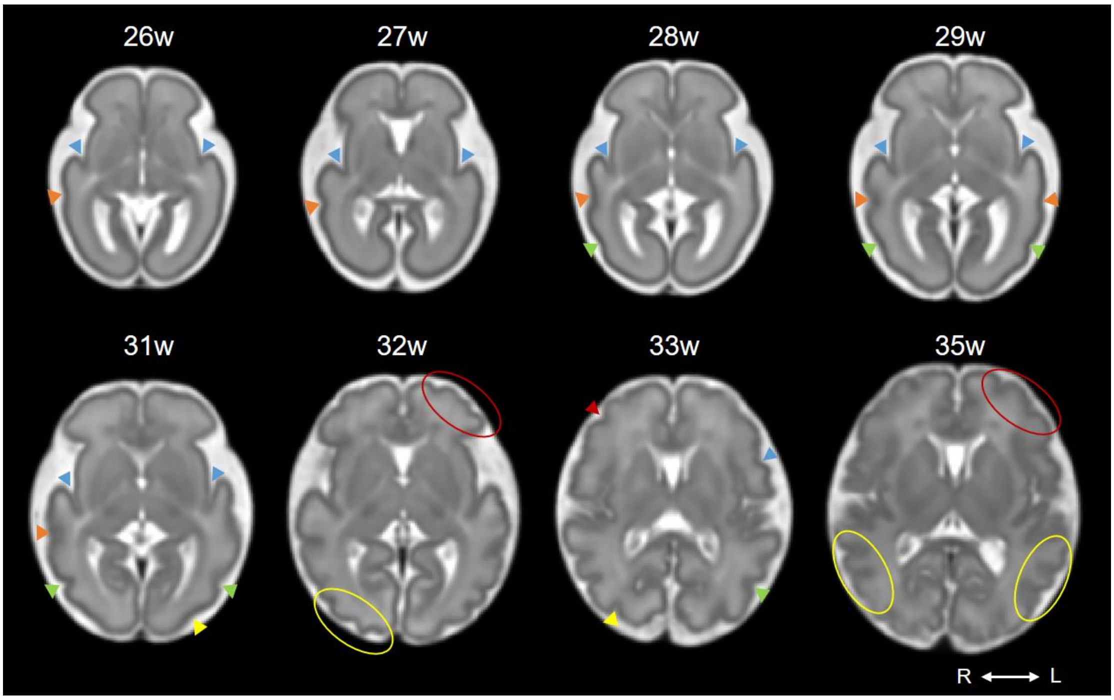


Fig. S3. Visual inspection of structural asymmetries of CHN atlas at representative GA. Arrows and circles show the asymmetrical sulci and gyri.

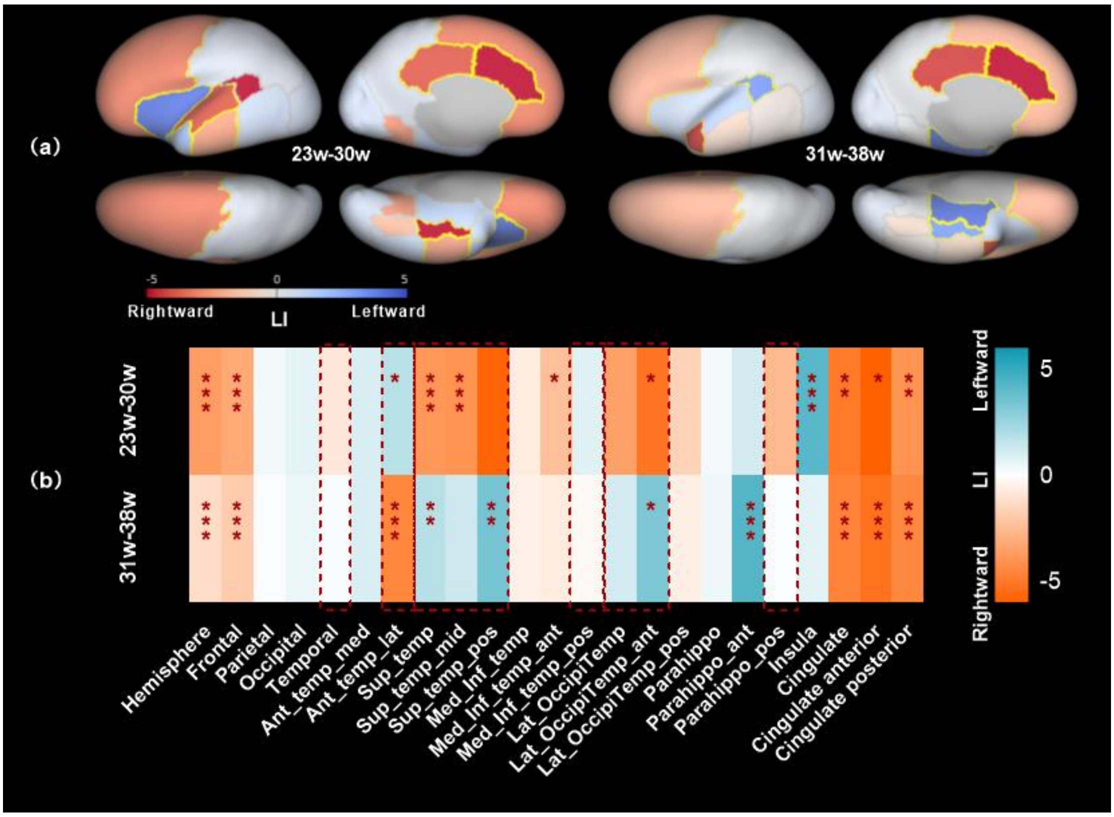


Fig. S4. Cortical asymmetry in terms of curvature. (a) LI maps of the fetal brain in the 23-30w and 31-38w groups. Positive LI indicates leftward asymmetry and negative LI indicates rightward asymmetry. Significant asymmetric regions are indicated by yellow outlines. (b) Statistical analysis of the LIs in the two GA groups. Red dashed box indicated the regions in which lateralization reversed between the two

GA groups. *Adjusted P <0.05, **adjusted P<0.01, ***adjusted P <0.001. Abbreviations: Sup=Superior, Temp=Temporal, Med=Medial, Inf=Inferior, Lat=Lateral, pos=posterior, ant=anterior, Occipitemp=Occipitotemporal, Parahippo=parahipppocampal.

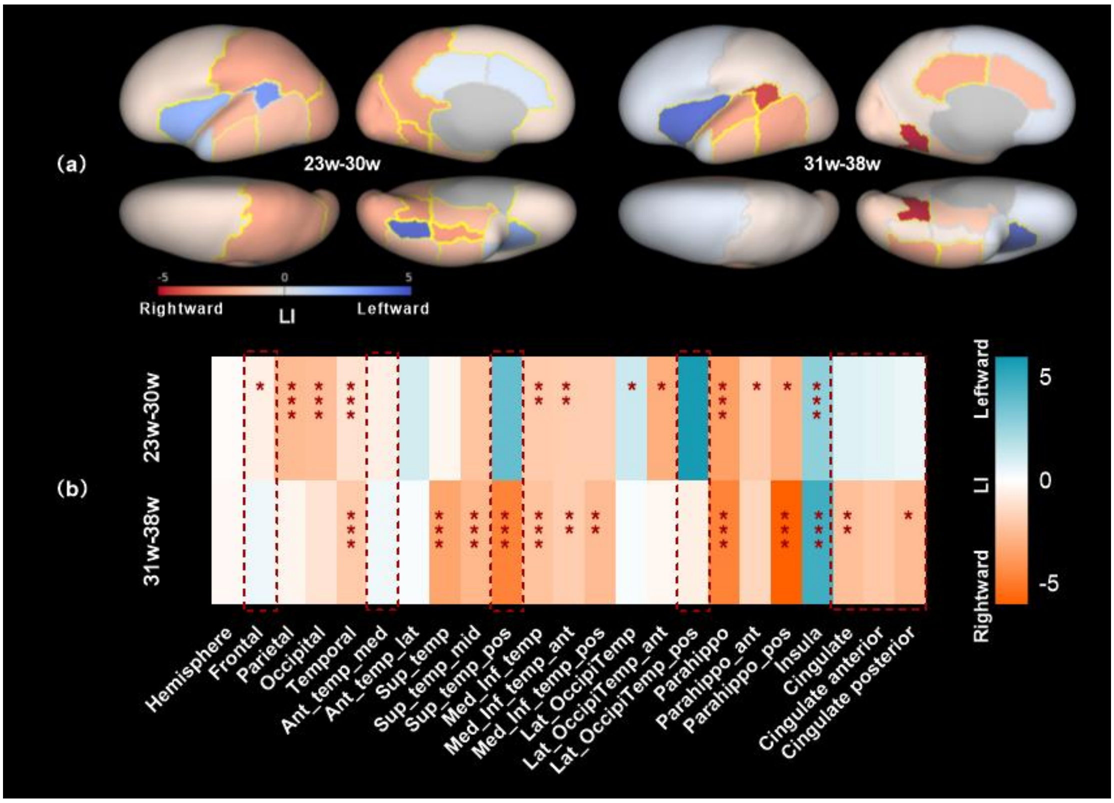


Fig. S5. Cortical asymmetry in terms of sulcal depth. (a) LI maps of the fetal brain in the 23-30w and 31-38w groups. Positive LI indicates leftward asymmetry and negative LI indicates rightward asymmetry. Significant asymmetric regions are indicated by yellow outlines. (b) Statistical analysis of the LIs in the two GA groups. Red dashed box indicated the regions in which lateralization reversed between the two GA groups. *Adjusted P <0.05, **adjusted P<0.01, ***adjusted P <0.001. Abbreviations: Sup=Superior, Temp=Temporal, Med=Medial, Inf=Inferior, Lat=Lateral, pos=posterior, ant=anterior, Occipitemp=Occipitotemporal, Parahippo=parahipppocampal.

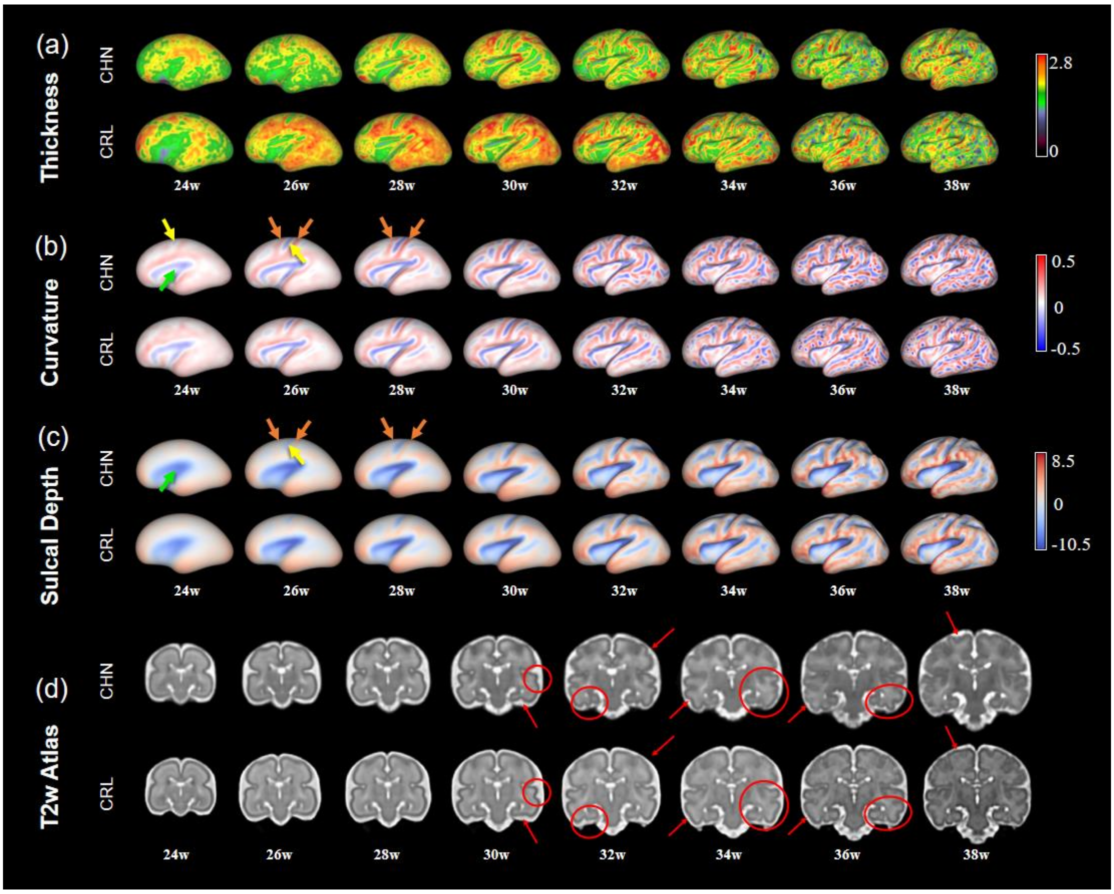


Fig. S6. Comparison of our proposed CHN fetal brain atlas and the CRL atlas at representative GAs. Vertex-wise cortical morphological indices of CHN atlas and CRL atlas, including (a) thickness, (b) curvature, and (c) sulcal depth, on the inflated surface. (d) Coronal views of the two atlases. The primary differences were indicated by red circles and arrows showing that CHN atlas had more convoluted gyrification in temporal lobe. Green arrows: Sylvian fissure; yellow arrows: central sulcus; orange arrows: precentral and postcentral gyri.

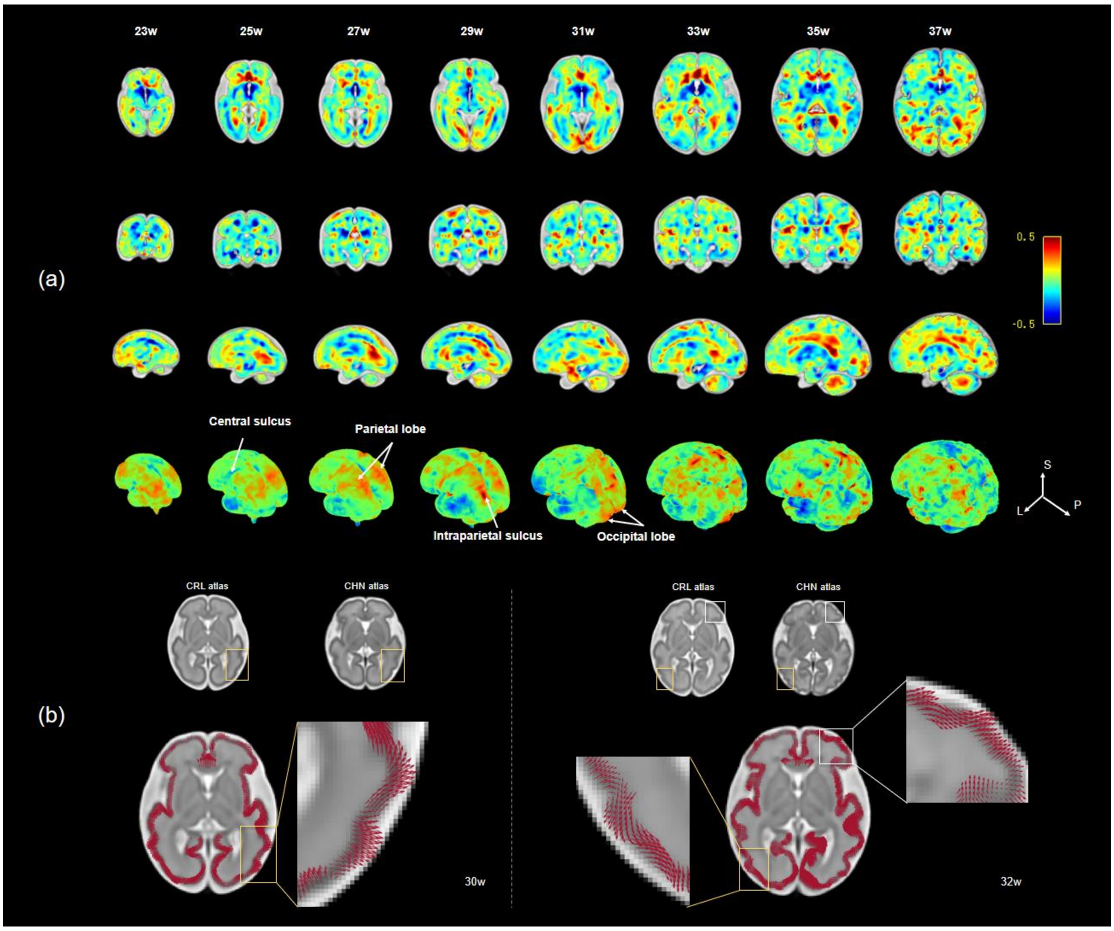


Fig. S7. (a) Morphological differences between CHN and CRL atlas based on the logarithm of the Jacobian determinant from SyN registration by transforming CHN atlas to CRL atlas at representative GAs. The colors in red and yellow represent local expansion, indicating more convex gyri in our CHN atlas than that in the CRL atlas, and green and blue show local shrinkage, indicating deeper sulci in our atlas than that in the CRL atlas. (b) TBM-based deformation fields at 30w and 32w GA. The growth vector indicated the direction of the morphological difference in CHN atlas compared to CRL atlas. For example, the growth vectors on gyri pointed outward and the growth vectors on sulci pointed inward indicating more convex gyri and deeper sulci in the CHN brains.
